# Targeting IL-1/IRAK1/4 signaling in Acute Myeloid Leukemia Stem Cells Following Treatment and Relapse

**DOI:** 10.1101/2024.11.09.622796

**Authors:** Tzu-Chieh Ho, Mark W. LaMere, Hiroki Kawano, Daniel K. Byun, Elizabeth A. LaMere, Yu-Chiao Chiu, Chunmo Chen, Jian Wang, Nikolay V. Dokholyan, Laura M. Calvi, Jane L. Liesveld, Craig T. Jordan, Reuben Kapur, Rakesh K. Singh, Michael W. Becker

## Abstract

Therapies for acute myeloid leukemia (AML) face formidable challenges due to relapse, often driven by leukemia stem cells (LSCs). Strategies targeting LSCs hold promise for enhancing outcomes, yet paired comparisons of functionally defined LSCs at diagnosis and relapse remain underexplored. We present transcriptome analyses of functionally defined LSC populations at diagnosis and relapse, revealing significant alterations in IL-1 signaling. Interleukin-1 receptor type I (IL1R1) and interleukin-1 receptor accessory protein (IL1RAP) were notably upregulated in leukemia stem and progenitor cells at both diagnosis and relapse. Knockdown of *IL1R1* and *IL1RAP* reduced the clonogenicity and/or engraftment of primary human AML cells. In leukemic MLL-AF9 mice, *Il1r1* knockout reduced LSC frequency and extended survival. To target IL-1 signaling at both diagnosis and relapse, we developed UR241-2, a novel interleukin-1 receptor-associated kinase 1 and 4 (IRAK1/4) inhibitor. UR241-2 robustly suppressed IL-1/IRAK1/4 signaling, including NF-κB activation and phosphorylation of p65 and p38, following IL-1 stimulation. UR241-2 selectively inhibited LSC clonogenicity in primary human AML cells at both diagnosis and relapse, while sparing normal hematopoietic stem and progenitor cells. It also reduced AML engraftment in leukemic mice. Our findings highlight the therapeutic potential of UR241-2 in targeting IL-1/IRAK1/4 signaling to eradicate LSCs and improve AML outcomes.

## Introduction

Acute myeloid leukemia (AML) is a hematological malignancy predominantly affecting older adults, with an annual incidence rate of 2 to 4 cases per 100,000 individuals (1). The standard induction chemotherapy regimen for AML, known as the “7+3” protocol, involves a combination of cytarabine for 7 days and anthracyclines for 3 days (2). Despite these treatments, AML patients still face high relapse rates or succumb to the disease. Young patients (aged <60 years) report 5-year survival rates of 30-35%, while older patients (aged ≥60 years) experience 5-year survival rates of less than 10-15% (3). It is widely acknowledged that primitive leukemia stem cells (LSCs) and pre-leukemic populations sustain AML and drive its recurrence (4). We previously assessed the evolution of LSC population(s) during patients’ clinical courses and identified a 9-90-fold increase in LSC frequency and a notable expansion in phenotypic diversity of the LSC population (5). Despite ongoing efforts to develop novel therapies targeting malignant hematopoietic stem and progenitor (HSPC) populations, current treatments for relapsed AML remain limited and largely ineffective for the majority of patients (6-9). Consequently, there exists a pressing need for therapies that specifically target LSC populations across the entire disease spectrum, with the potential to improve overall survival (OS) rates for AML patients.

Hematopoietic disorders associated to aging, clonal hematopoiesis, myelodysplastic syndromes (MDS), and AML have been linked to the dysregulation of innate immune, inflammatory pathways, as well as systemic inflammation (10, 11). Interleukin-1 (IL-1)/Toll-Like Receptor (TLR) signaling pathways are potent inflammatory cascades that respond to infections and various stimuli (12, 13). They can induce multiple critical downstream mechanisms involved in gene regulation and cell survival, including NF-κB, p38, JNK, and AP-1, mediated by MyD88 adaptor and interleukin-1 receptor-associated kinase 1 and 4 (IRAK1/4) (14). Chronic exposure to IL-1 can impede the self-renewal capacity of HSCs and promote myeloid differentiation; however, these effects are reversible upon IL-1 withdrawal (15). Recurrent genetic mutations observed in AML and MDS, such as spliceosomes and epigenetic modifier mutations, have been linked to alterations in IL-1/TLR signaling (16-18). Furthermore, studies, including our own, have demonstrated overexpression of key components of IL-1/TLR signaling, including the IL-1 ligand, receptor, and coreceptor, in AML and MDS patients, suggesting their potential as therapeutic targets for these diseases (5, 19, 20). Nonetheless, it remains uncertain whether these aberrant alterations in IL-1/TLR signaling sustain their effects on AML LSCs following treatment and during relapse.

Several agents, either approved for clinical use or undergoing investigation in clinical trials, have been developed for targeting IL-1/TLR signaling. Anakinra, a recombinant human interleukin-1 receptor antagonist (rhIL1-Ra), is commonly used in clinical settings to manage IL-1 driven autoimmune and autoinflammatory diseases, such as rheumatoid arthritis (RA) (21, 22). IRAK4 inhibitors, such as PF06650833 (Zimlovisertib) and CA-4948 (Emavusertib), have been or are currently being evaluated in clinical trials for RA (NCT02996500) and myeloid malignancies (NCT04278768), respectively (23). Furthermore, proteolysis targeting chimeric (PROTAC) degraders targeting IRAK1/4 proteins are being evaluated in pre-clinical and clinical trials for hematologic malignancies, including non-Hodgkin lymphoma (23). Nevertheless, the efficacy of compounds targeting IL-1/TLR/IRAK1/4 signaling in eradicating LSCs in AML patients at various disease stages, such as diagnosis, remission, and relapse, remains to be determined.

In this study, we have uncovered that upregulated IL-1 signaling may be a mechanism underlying changes in functionally defined LSC populations observed in paired samples from AML patients at diagnosis and relapse. Loss of IL-1 signaling factors, including IL1RAP and IL1R1, significantly affects LSC function *in vitro* and *in vivo*. To counteract IL-1 signaling, we developed a novel small molecule compound targeting IRAK1/4 (UR241-2). UR241-2 effectively suppresses IL-1/IRAK1/4 downstream signaling, including the activation of NF-κB and the phosphorylation of P65 and P38, consequently impeding the LSC clonogenicity and engraftment capability. UR241-2 also suppresses LSC function in primary human AML cells at diagnosis and in relapse, with minimal impact on normal bone marrow (NBM) HSPCs. Overall, our findings suggest that targeting IL-1/IRAK1/4 signaling with UR241-2 represents a promising therapeutic strategy to eradicate LSCs across different disease stages, potentially leading to improved outcomes for AML patients.

## Material and methods

Supplemental Methods are available on the *bioRxiv* website in the Supplemental Data file for additional information and details.

### Primary human samples

Eighteen diagnostic and relapsed AML samples were obtained and characterized as previously described (5). We also included nine other AML specimens for immunophenotyping or functional assays (supplemental Table 1). Most of these patients were treated with 7+3 regimen. Bone marrow (BM) and peripheral blood (PB) samples from AML patients or normal donors were obtained following informed consent and in accordance with protocols approved by the institutional review board (IRB) at the University of Rochester Medical Center (URMC) and Roswell Park Comprehensive Cancer Center. All samples were processed following the methods outlined in our previous work (5).

### TaqMan Low Density Array (TLDA) assay

Total RNA was extracted from human LSC and HSC populations using TRIzol Reagent (Invitrogen) and cleaned-up using RNeasy column (Qiagen). The cDNA sample was generated using the iScript cDNA Synthesis Kit (Bio-Rad) and pre-amplified using pooled TaqMan Gene Expression Assay primer/probe combinations and TaqMan PreAmp Master Mix (Thermo Fisher Scientific). Custom TLDA cards were designed with 157 genes selected from our previous work (24) and other LSC references (25) (supplemental Table 2). Our previous work identified 3,005 differentially expressed genes (DEGs) and 10 pathways dysregulated in enriched phenotypically defined LSC populations (CD34^+^CD38^-^CD123^+^Lin^-^ or CD34^+^CD38^-^CD90^-^Lin^-^) compared to normal HSC populations (CD34^+^CD38^-^CD90^+^Lin^-^) using genome-wide expression analyses and two independent pathways analyses (24). For this study, we selected 124 of the 3005 DEGs based on their involvement in five dysregulated pathways (Adherens Junction, Wnt, Regulation of Actin Cytoskeleton, Jak-STAT, and MAPK) or reported roles in response to therapy. Additionally, 27 genes were selected from a 42 gene LSC-related (LSC-R) gene signature reported by *Eppert et al*. (25) in which they compared the expression profiles of functionally defined LSC populations to non-LSC populations independent of surface antigen phenotype. Ten of these 27 genes overlapped with our 3,005 DEG list. RT-qPCR was carried out using the 7900HT Sequence Detection System (Applied Biosystems) and analyzed using ExpressionSuite Software (Applied Biosystems), with *GAPDH* and *HPRT1* used for normalization. Gene expression levels generated from independent runs were normalized to universal human reference RNA (UHRR) standard (Stratagene), a positive control composed of total RNA from 10 human cell lines used as a constant reference in gene-profiling analyses.

### NSG xenotransplantation assay

To conduct shRNA knockdown xenograft studies, young NOD.Cg-*Prkdc^scid^ Il2rg^tm1Wjl^*/SzJ (NSG) mice were irradiated with a sublethal dose (250 cGy) 24 hours prior to transplantation. Recipient mice also received two doses of 0.5 mg/g of human intravenous immune globulin (IVIG) through intraperitoneal (IP) injection on the day of irradiation and on the day of transplantation. Live T-cell-deplete GFP^+^ primary AML cells (DAPI^-^GFP^+^CD3^-^) were sorted and transplanted into recipient mice via the tail-vein route in a final volume of 0.2 ml of 1x PBS plus 0.5% FBS. Equal numbers of sorted cells from each group (shRNA knockdown or scramble control groups) were transplanted into recipients. Populations with fewer than one million cells per mouse were mixed with syngeneic NSG splenocytes before transplantation. Mice were sacrificed 12 weeks post transplantation or when they appeared moribund. Leukemic burden was defined by the presence of live GFP^+^ cells in mouse BM using flow cytometry. Recipient BM cells were also stained with antibodies specific to mouse CD45, human CD45, human CD19, and human CD97 to determine the tumor burden, as previously described (5).

### Limiting dilution analysis (LDA)

The limiting dilution analysis was performed as previously described (26). In brief, BM cells from primary transplantations of MLL-AF9 *Il1r1* wild type (IL1R1 WT) and MLL-AF9 *Il1r1* knockout (*Il1r1*^-/-^; IL1R1 KO) mice (three mice per group) were pooled and serially diluted for secondary transplantations to achieve doses of 5, 50, 500, 5,000, and 50,000 cells per mouse. Prior to transplantation, the test cells were mixed with fresh CD45.1 BM competitor cells to achieve 1 × 10^6^ cells per mouse. These cells were then injected into lethally irradiated Pep Boy recipient mice (5 Gy × 2) via the tail-vein route. Mice were monitored for leukemia signs throughout the study, and the frequency of LSCs was calculated using LCalc Software (STEMCELL Technologies).

### Synthesis of UR241-2

6-(5-Pyrazolyl)pyridine-2-carboxylic acid (0.1 mM, Accela ChemBio Inc.) in dry *N,N*-Dimethylformamide (DMF) (Sigma Aldrich) was dissolved and stirred with *N,N’*-Dicyclohexylcarbodiimide (DCC) (0.011 mM, Sigma Aldrich) in a round bottom flask covered with a drying tube, and placed in an ice bath for 15-20 minutes. Subsequently, the mixture was added to 4-(4-amino-3-methoxyphenyl)-1lambda6-thiomorpholine-1,1-dione hydrochloride (0.01 mM, Enamine) and stirred overnight. DMF was removed under reduced pressure, and the crude product was suspended in ethyl acetate, then washed successively with water and saline solution. The organic layer was separated, dried over anhydrous sodium sulfate, and filtered. Ethyl acetate was evaporated under pressure, yielding the crude product, which was subsequently purified using a preparative thin layer chromatography plate with DCM/MeOH (95:5) as eluent. The band containing the desired product was collected, and the compound was eluted from the silica gel using MeOH:DCM (9:1) as solvent system. The solvent was removed using a rotary evaporator, and UR241-2 was obtained as a yellowish powder after drying under vacuum. The compound was stored at -20^°^C. Mass spectrometry (MS) analysis yielded an expected mass of 427, with an observed mass of 428.

### In silico docking

The protein structure prediction tool Swiss-Model, based on homology modeling, was utilized to predict the structures of IRAK1 and IRAK4. Although crystal structures were available for these proteins, Swiss-Model was employed primarily to fill in missing residues and atoms. Specifically, IRAK1 was modeled using 6BFN as the template, while IRAK4 utilized 6UYA as the template. Subsequently, the modeled structures underwent optimization with Chiron. MedusaDock (27, 28) was employed to dock PF06650833 and UR241-2 to IRAK1 and IRAK4. The evaluation of docking poses relied on MedusaScore (29), a scoring function based on a custom physics-based force field. Each protein-ligand pair underwent 1000 docking attempts, and resulting docking poses were clustered by MedusaDock to select the centroid as the final candidate.

### Kinase profiling assay

The kinase profiling assay for UR241-2 involved testing in 10-dose IC_50_ mode with a 3-fold serial dilution, starting at 20 µM. The control compound, staurosporine, was tested in 10-dose IC_50_ mode with a 4-fold serial dilution, starting at 20 µM. Reactions were carried out with 1 µM ATP, and the % enzyme activity was calculated relative to DMSO controls. Curve fits were performed where the enzyme activities at the highest concentration of compounds were less than 65%. An IC_50_ value of less than 1.0 nM was estimated based on the best available curve fitting.

### Kinome map analysis

To perform kinome map analysis, UR241-2 was tested against 682 kinases in single-dose duplicate mode at a concentration of 5 nM. As a control compound, staurosporine was also tested in 10-dose IC_50_ mode with 4-fold serial dilution, starting at either 20 or 100 µM. Additionally, alternate control compounds (LDN193189, GW5074, D4476, Ro-31-8220, SCH772984, JNKi VIII, JNK-IN-7, PKR Inhibitor, SB202190, BI2536, Wee-1 Inhibitor, PI-103, NH125, and GSK-2606414) were tested in 10-dose IC_50_ mode with 3 or 4-fold serial dilution, starting at 10, 20, 50, or 100 µM. Reactions were carried out at 1 µM ATP, and the % enzyme activity was calculated relative to DMSO controls. The network diagram was generated using Rstudio “igraph” package. The kinome tree map was created using Coral generated from the Phanstiel Lab (30).

### NanoBRET Target Engagement Assay

HEK293 cells (RRID:CVCL_0045) were transiently transfected with 1 µg of IRAK1, IRAK4, FLT3, MAP4K1, MAP4K2, MAP4K3, MAP4K5, and LRRK2 NanoLuc Fusion vectors, along with 9 µg of transfection carrier DNA. Transfected cells were treated with UR241-2 and a reference compound for 1 hour using a 384-well assay format (4,000 cells/well). We used a starting concentration of 20 µM for UR241-2 and 10 µM for the reference compound, followed by 10 doses with 3-fold dilution. Target engagement was measured by NanoBRET assay. Curve fits were only performed when the % NanoBRET signal at the highest concentration of compounds was less than 55%.

### Drug treatment

To conduct drug treatment immunoblotting studies, THP-1 cells were cultured overnight in RPMI medium plus 0.5% FBS (1 × 10^6^ cells/mL). After serum-starvation, cells were pre-treated with UR241-2, PF06650833 (Selleck), and IRAK4-IN-7 (MCE) (4 µM) or vehicle (DMSO) for 30 minutes. Subsequently, human IL-1β (10 ng/mL) was added to the plates and incubated for 0, 10, and 30 minutes. At each time point, cells were collected, washed with cold 1x PBS, and lysed for immunoblotting assays. To perform drug treatment colony-forming unit cell (CFU-C) assays using murine cells, MLL-AF9 GFP^+^ splenocytes were sorted and mixed with IRAK1/4 inhibitors (UR241-2, PF06650833, and IRAK4-IN-7) at the indicated concentrations. The cells were then plated in methylcellulose-based medium for CFU-C assays (5 × 10^3^ cells/mL). Colonies were scored after 7 days of culture. To conduct drug treatment CFU-C assays using primary human cells, bulk MNCs from diagnostic and relapsed AML patients or NBM CD34^+^ cells sorted from three pooled donors were mixed with IRAK1/4 inhibitors (UR241-2, PF06650833, and IRAK4-IN-7) at the indicated concentrations. The cells were then plated in methylcellulose-based medium for CFU-C assays (1 × 10^4^ cells/mL for primary AML cells, 2.5 × 10^3^ cells/mL for NBM CD34^+^ cells). Colonies were scored after 14 days of culture.

### *Ex Vivo* treatment followed by transplantation

To conduct *ex vivo* treatment followed by transplantation, sorted MLL-AF9 GFP^+^ splenocytes were cultured in MLL-AF9 medium (IMDM supplemented with 10% FBS, 50 ng/mL recombinant murine SCF and Flt3-Ligand, 10 ng/mL IL-3 and IL-6 (all cytokines sourced from PeproTech), and 1x Pen-Strep) and treated with UR241-2 (4 µM), PF06650833 (4 µM), or vehicle (DMSO) for three days (5 × 10^5^ cells/0.5 mL per mouse). After treatment, cells were collected, resuspended in 1x PBS plus 0.5% FBS, and transplanted into sublethally irradiated (6.5 Gy) C57BL/6J mice through tail-vein injection (0.2 mL per mouse). Mice were sacrificed after four weeks to assess engraftment and leukemia burden.

### Sex as a biological variable

Sex was not considered as a biological variable.

### Statistics

The statistical analysis methods were specified in the figure legends. Statistical analyses were conducted using GraphPad Prism (GraphPad Software; RRID:SCR_002798). A *P*-value less than 0.05 was considered significant for all statistical analyses. Clustering and heat maps were generated using Multiple experiment Viewer (MeV). LSC and HSC data were clustered using the Pearson correlation metric with average linkage.

### Study approval

All samples from AML patients or normal donors were obtained with informed consent and in accordance with protocols approved by the IRB at the URMC (URCC: LEU 07047) and Roswell Park Comprehensive Cancer Center. Additionally, animal experiments conducted in this study were approved by the Institutional Animal Care and Use Committee (IACUC) at the URMC (UCAR-2005-256E).

### Data availability

All data generated by the present study are provided within the article and the Supplemental Data file. For original data, please contact corresponding authors.

## Results

### Identification of molecular signatures associated with LSC expansion after drug treatment and disease progression

In our previous work on examining the effects of therapy and disease progression in AML patients treated with conventional chemotherapy regimens, we demonstrated a notable increase in LSC frequency and expansion from diagnosis to relapse (5). We also observed genetic instability and alterations in surface and intracellular biomarker profiles at relapse. Previous studies have attempted to elucidate LSC signaling patterns through gene expression profiling of enriched LSC populations (24, 25, 31, 32). To delineate the impact of treatment and relapse on LSC signatures, we conducted targeted transcriptome profiling of paired, functionally defined LSC populations from AML patients at diagnosis and relapse using TaqMan Low Density Array (TLDA) cards, a RT-qPCR-based assay (Figure. 1A). Four distinct populations from AML patients were sorted based on CD34/CD38 or CD32/CD38 expression and the functionally defined LSC populations in these patients at both diagnosis and relapse were characterized using the NSG xenograft model (5). The LSC cells, functionally defined at both diagnosis and relapse, were sorted again from patient samples for the TLDA assay. Custom TLDA cards were designed with 157 genes selected from our previous work (24) and other LSC references (25) (supplemental Table 2). We observed significant inter-patient heterogeneity among AML populations, particularly notable for patients with LSCs defined by CD32 and CD38 (supplemental Figure 1A). Hierarchical clustering demonstrated that LSC populations from individual patients tended to cluster together rather than diagnosis or relapse samples from different patients (supplemental Figure 1A). Comparison of paired sample data revealed 8 upregulated and 28 downregulated genes in relapse samples (paired *t*-test *P* < 0.05) (supplemental Figure 1B). The upregulated genes are associated with several pathways, including Jak-STAT, Apoptosis, Tight junction, Adherens, ARENRF2, and others. Gene Set Enrichment Analysis (GSEA) (33) indicated positive associations between relapse and LSC activities: the upregulated (or downregulated) genes were highly expressed in functionally defined LSC^+^ (or LSC^-^) cell fractions of two published xenotransplantation models (25, 34) (*P*-values < 0.05; Figure 1B and supplemental Figure 1C). *ATP1B1*, *IL1RAP*, *IL3RA*, and *WT1* were found in both leading-edge components (Figure 1B), suggesting their pivotal roles in LSCs and AML relapse. Interleukin-1 receptor accessory protein (IL1RAP) is a coreceptor of interleukin-1 receptor type I (IL1R1), capable of responding to extrinsic IL-1 and initiating downstream signaling (35). Taken together, our findings highlight molecular signatures associated with LSC expansion following drug treatment and disease progression, notably including IL-1 signaling. However, it is unclear if IL-1 signaling is functionally relevant in relapsed LSCs versus LSCs at diagnosis.

**Figure 1.**
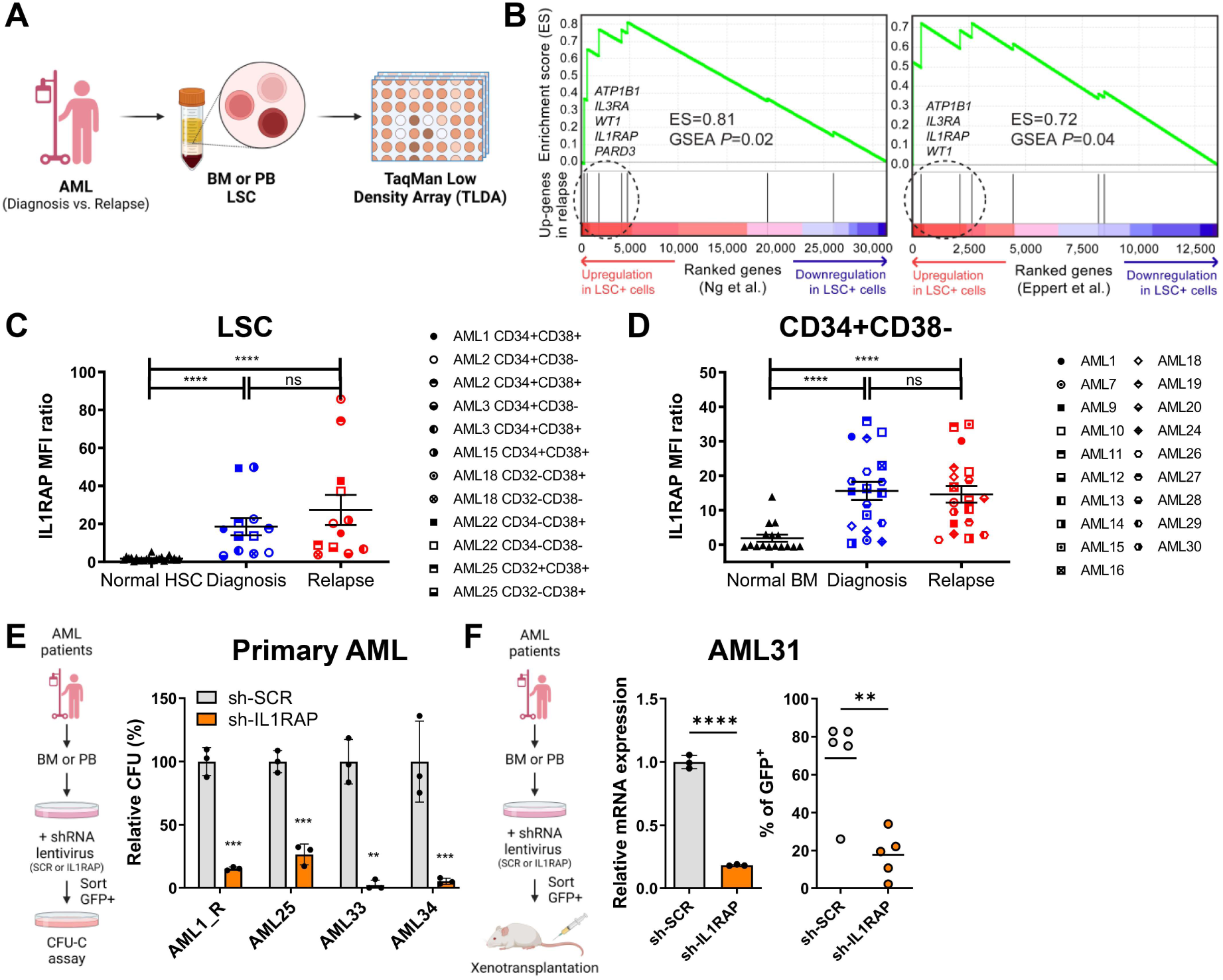
IL1RAP is upregulated in functionally defined LSC populations at diagnosis and relapse in AML. (A) Schematic of targeted transcriptome analyses of paired AML diagnostic and relapsed LSC populations. LSC populations were functionally defined using the NOD.Cg-*Prkdc^scid^ Il2rg^tm1Wjl^/SzJ* (NSG) model, as described in *Ho et al.(5)*. Briefly, AML cells were sorted based on CD3^-^CD34/CD38 or CD3^-^CD32/CD38 into four populations and transplanted into NSG mice to assess their engraftment ability. The functionally defined LSC phenotypes vary among patients. Subsequently, AML LSC populations were sorted from bone marrow (BM) or peripheral blood (PB) specimens, and the gene expression was analyzed using TaqMan Low Density Array (TLDA) assay. Twelve paired AML LSC populations (CD3^-^CD34/CD38 or CD3^-^CD32/CD38) from seven samples and four NBM HSC populations (CD34^+^CD38^-^CD90^+^Lin^-^) were analyzed by TLDA-based analyses. (B) Gene Set Enrichment Analysis (GSEA) plots comparing genes upregulated upon relapse with published expressional profiles of LSC^+^ cells. Genes were ranked based on the significance of differential expression associated with LSC^+^ cells (indicated by red and indigo arrows) of two published datasets, *Eppert et al.*(25) and *Ng et al.*(34). Eight upregulated genes were defined as a gene set (black segments) and tested for the enrichment (enrichment score; green curve) at either side of the ranked list. Denoted by a dashed circle are leading-edge component genes that contribute to the enrichment. Significance was assessed by the random permutation test. WT1 was analyzed in six pairs of samples. (C) Protein expression levels of IL1RAP on primary AML LSCs at diagnosis and relapse compared to NBM HSCs (n = 24, CD34^+^CD38^-^). Median fluorescence intensity (MFI) of the cell-surface IL1RAP was quantified by flow cytometry analysis and normalized to Fluorescence Minus One (FMO) control. MFI ratio = (Sample MFI – FMO MFI)/FMO MFI. Data are shown as Mean ± standard error of the mean (SEM). The two-tailed Mann-Whitney test (NBM vs. AML) or Wilcoxon matched-pairs signed rank test (diagnosis vs. relapse) were used in comparison. (D) Protein expression levels of IL1RAP on primary AML CD34^+^CD38^-^ populations at diagnosis and relapse compared to NBM counterparts (n = 16, CD34^+^CD38^-^). The MFI ratio and statistical analyses were conducted as described in *(C)*. (E) Colony-forming unit cell (CFU-C) assays of primary AML cells with or without knockdown of IL1RAP (sh-SCR: scramble shRNA; sh-IL1RAP: IL1RAP shRNA). Sorted GFP^+^ cells (AML1, AML34) or GFP^+^CD3^-^ cells (AML25 and AML33) were plated post lentiviral infection, and CFU-C was counted after culture. AML, n = 4. Mean ± standard deviation (SD). Unpaired t test. (F) Xenotransplantation assays of primary AML cells (AML31) with or without knockdown of IL1RAP. (Left panel) Relative gene expression levels of *IL1RAP* in AML31 with or without knockdown of IL1RAP. Total RNA was isolated from sorted GFP^+^CD3^-^ cells post infection. Mean ± SD. Unpaired t test. (Right panel) Each dot represents the GFP^+^ engraftment level in BM of individual NSG mouse analyzed by flow cytometry analysis. The horizontal line is the mean of GFP^+^ engraftment level in NSG mice of each group. Unpaired t test. \*\**P* ≤ 0.01, \*\*\**P* ≤ 0.001, \*\*\*\**P* ≤ 0.0001, ns = not significant. Schematics were created with BioRender.com.

### IL1RAP is upregulated in LSCs at diagnosis and in relapse

The IL-1 family comprises 11 cytokines, including IL-1α and IL-1β, which exert comparable biological activities and are potent regulators of inflammation (12, 36). Upon binding to the IL-1R and its coreceptor IL1RAP, IL-1 initiates various several downstream pathways associated with both innate and adaptive immunity (12, 14). Previous reports have demonstrated the overexpression of IL1RAP on the surface of cells from chronic myeloid leukemia (CML), AML, and MDS relative to their normal counterparts (19, 37-39). However, the impact of IL1RAP expression in AML following treatment and disease progression remains unclear. To address a potential role for IL1RAP in relapse, we investigated cell surface IL1RAP expression in paired functionally defined LSC populations and observed a significant increase of IL1RAP in LSCs at diagnosis from most AML patients compared to NBM HSCs (CD34^+^CD38^-^ cells) (Figure 1C). Notably, IL1RAP expression levels in NBM HSCs were consistently low (Figure 1C), consistent with previous findings (5, 19, 37). Importantly, IL1RAP was consistently overexpressed in LSCs at relapse (NBM HSCs = 1.7±1.1, diagnostic AML LSCs = 3.2 to 49.9, relapsed AML LSCs = 3.8 to 85.7) (Figure 1C). We further demonstrated that IL1RAP expression in leukemia stem and progenitor cells (LSPCs) CD34^+^CD38^-^ populations was upregulated in paired diagnostic and relapsed AML samples compared to their NBM counterparts (Figure 1D). Similar observations were made in paired primary human AML bulk, CD34^+^ and CD34^+^CD38^+^ populations (supplemental Figure 2A-C).

Previous studies demonstrate that elevated *IL1RAP* expression levels are independently associated with poor OS in AML with normal karyotype (19). To further explore the functional significance of IL1RAP in AML LSC populations at both diagnosis and relapse, we targeted *IL1RAP* genes in human primary AML cells and leukemia cell lines using short-hairpin RNA (shRNA) lentiviruses (supplemental Figure 3A-B). Our results revealed that loss of IL1RAP significantly reduces short-term clonogenicity in human primary AML cells (Figure 1E) and the U937 leukemia cell line (supplemental Figure 3C), consistent with previous reports (19, 37). To evaluate the long-term functional effects of IL1RAP loss in LSCs, xenotransplantation assays were conducted using NOD/SCID/IL2rγ^-/-^ (NSG) mice as recipients for infected cells. Knockdown of IL1RAP via shRNA generally impeded leukemia burden in recipient mice, underscoring the crucial functional roles of IL1RAP in stem cell populations in primary AML (Figure 1F and supplemental Figure 3D). These findings provide novel insights regarding the role of IL-1 signaling in regulating the growth and survival of LSCs both at diagnosis and at relapse.

### IL-1 signaling plays an important role in LSCs at diagnosis and relapse

IL-1 induces the activation of several critical downstream pathways, such as nuclear factor κB (NF-κB), mitogen-activated protein kinases/extracellular signal-regulated kinases (MAPK/ERK), and c-Jun N-terminal kinase (JNK) (Figure 2A). Recent reports have highlighted the upregulation of IL-1β and IL1R1 levels in AML patients (20). Interleukin 1 Receptor Associated Kinase 3 (IRAK3), a member of the IRAK protein family, serves as a key negative regulator in IL-1/TLR signaling (40). Our TLDA analysis revealed a significant downregulation of *IRAK3* in functionally defined LSC populations from diagnosis to relapse in AML patients (Figure 2B), suggesting a decrease in negative regulation on IL-1 signaling following treatment and disease progression.

**Figure 2.**
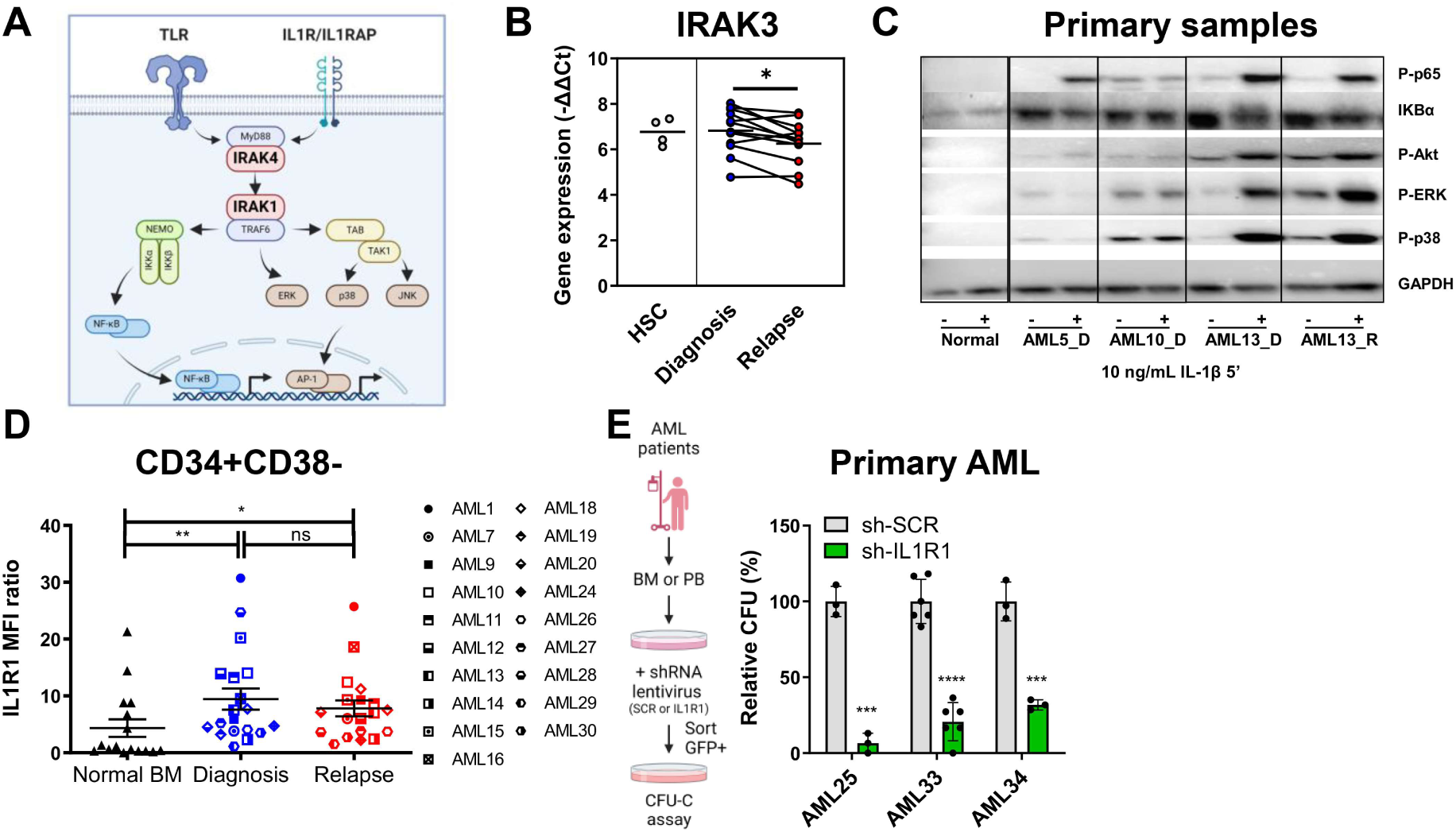
IL-1 signaling plays an important role in LSCs at diagnosis and in relapse. (A) Schematic of IL-1/TLR signaling. (B) Gene expression levels of IRAK3 (IRAKM) in AML LSCs from diagnostic and relapsed samples as well as NBM HSCs. Samples were processed and analyzed using TLDA assay as described in Figure 1A. AML LSC, n = 12. NBM HSC, n = 4. The horizontal line is the mean of gene expression levels. Two-tailed Wilcoxon matched-pairs signed rank test (diagnosis vs. relapse). (C) Immunoblotting analyses of IL-1 signaling factors in primary AML and NBM samples, cultured with or without IL-1β treatment (10 ng/mL for 5 minutes). (D) Protein expression levels of IL1R1 in CD34^+^CD38^-^ populations of NBM (n = 16) or AML diagnostic and relapsed samples (n = 19). The MFI ratio and statistical analyses were conducted as described in Figure 1C. (E) CFU-C assays of primary AML cells with or without knockdown of IL1R1 (scramble shRNA: sh-SCR; IL1R1 shRNA: sh-IL1R1). Mean ± SD. Unpaired t test. \**P* ≤ 0.05, \*\**P* ≤ 0.01, \*\*\**P* ≤ 0.001, \*\*\*\**P* ≤ 0.0001, ns = not significant. Schematics were created with BioRender.com.

To further investigate IL-1-mediated signaling, we performed immunoblotting of primary human AML cells stimulated with IL-1β. These studies demonstrated the activation of IL-1 signaling in both diagnostic and relapsed AML samples, evidenced by increased phosphorylation of p65 and p38 (Figure 2C). Conversely, IL-1 signaling in NBM cells following IL-1 stimulation exhibited lesser impacts (Figure 2C), potentially correlated with the lower levels of IL1RAP/IL1R1 on NBM cells. Further validation of our findings was performed using leukemia cell lines (KG-1a, THP-1, and U937) incubated with IL-1β. IL-1β treatment revealed increased phosphorylation of p65 and p38 within a short timeframe (5-10 minutes) (supplemental Figure 4A-C). Additionally, the phosphorylation of ERK and JNK exhibited variable activation patterns in these cells (supplemental Figure 4A-C). Overall, our findings confirm the activation of IL-1 signaling in leukemia cells following IL-1β stimulation at both diagnosis and relapse.

Our observations revealed a significant upregulation of IL1R1 in the majority of AML CD34^+^CD38^-^ cells at diagnosis and relapse, compared to NBM CD34^+^CD38^-^ cells (Figure 2D). Interestingly, IL1R1 expression did not increase in AML bulk, CD34^+^, and CD34^+^CD38^+^ populations at diagnosis and relapse compared to their normal counterparts (supplemental Figure 5A-C). To determine if IL1R1 inhibition regulates the growth and survival of LSCs, we employed shRNA lentiviruses to knock down IL1R1 in human primary AML samples and leukemia cell lines (supplemental Figure 6A-B). Our findings demonstrated that loss of IL1R1 impaired the clonogenicity of human primary AML cells (Figure 2E) and leukemia THP-1 cell line (supplemental Figure 6C). No reduction in colony-forming units was observed when IL1R1 was knocked down in NBM CD34^+^ cells (supplemental Figure 6D).

### Loss of IL1R1 reduces LSC frequency and prolongs overall survival in MLL-AF9 model

To investigate the potential role of IL1R1 in leukemia progression, we utilized a leukemic murine model transformed by the *MLL-AF9* oncogene (41) and assessed the impact of IL1R1 depletion on LSC frequency *in vivo*. Lin^-^Sca-1^+^c-Kit^+^ (LSK) BM cells from wild type (WT; IL1R1 WT) and *Il1r1* knockout (*Il1r1*^-/-^; IL1R1 KO) mice were isolated and infected with a retrovirus co-expressing the MLL-AF9 fusion product and green fluorescence protein (pMSCV-MLL-AF9-GFP). Infected cells were subsequently transplanted into B6.SJL-*Ptprca Pepcb*/BoyJ (Pep Boy) recipients. BM cells harvested from primary MLL-AF9 IL1R1 WT and MLL-AF9 IL1R1 KO recipients were diluted and transplanted into secondary recipients (5 to 50,000 cells per mouse) (Figure 3A). All mice that received 50,000 cells died within 31 days (Figure 3B). However, depletion of IL1R1 significantly delayed the time to death of mice receiving fewer than 5,000 cells (IL1R1 WT: 28-140 days, IL1R1 KO: 40-140 days. *P*-value < 0.0001. Data collection stopped on Day 140) (Figure 3B). Limiting-dilution calculations show the LSC frequency was 1:152 in IL1R1 KO mice, which was five-fold lower than the 1:29 in IL1R1 WT mice (Figure 3C). Altogether, our findings suggest that IL1R1 plays an important role in maintaining LSC functions. Depletion of IL1R1 can decrease LSC frequency and prolong OS in AML.

**Figure 3.**
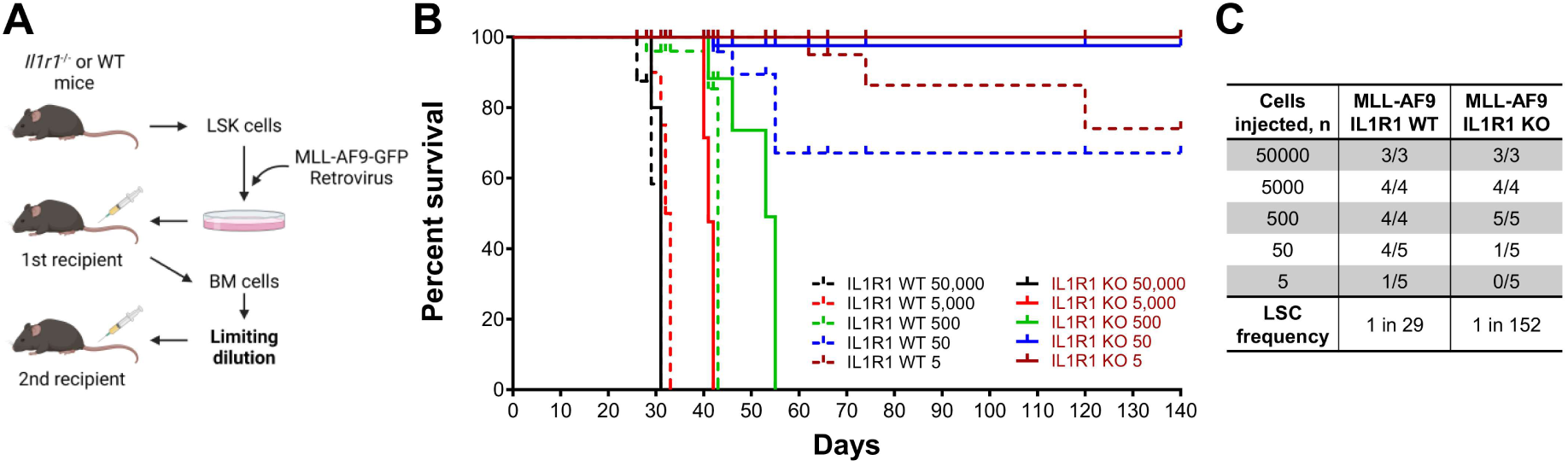
Loss of IL1R1 reduces LSC frequency and prolongs overall survival in MLL-AF9 model. (A) Schematic of the experiment strategy for limiting dilution analysis (LDA) in leukemia MLL-AF9 mice. In brief, LSK cells (Lin^-^Sca-1^+^c-Kit^+^) from wide type (WT; IL1R1 WT) and *Il1r1* knockout (*Il1r1*^-/-^; IL1R1 KO) mice were sorted and infected with a retrovirus co-expressing the MLL-AF9 fusion product and green fluorescence protein (MLL-AF9-GFP). GFP^+^ BM cells were sorted from primary recipients and limiting dilution cells were transplanted into secondary Pep Boy recipients (5 to 50,000 cells per mouse with competitor CD45.1 BM cells). Secondary recipients were followed until time of death. (B) Overall survival analyses of recipients transplanted with limiting dilution cells (5 to 50,000) with MLL-AF9 IL1R1 WT and MLL-AF9 IL1R1 KO as described in *(A)*. (C) The LSC frequencies in MLL-AF9 IL1R1 WT and MLL-AF9 IL1R1 KO mice transplanted with 5 to 50,000 cells as described in *(A)*. Schematics were created with BioRender.com.

To determine which LSPC populations are impaired in the absence of IL1R1, IL1R1 WT and IL1R1 KO MLL-AF9-GFP splenocytes from primary recipients were sorted for leukemia LSK, CMP, and GMP populations (LSK: Lin^-^Sca-1^+^c-Kit^+^; CMP: Lin^-^c-Kit^+^(16/32)FCγR^-^CD34^+^; GMP: Lin^-^c-Kit^+^(16/32)FCγR^+^CD34^+^) and then transplanted into secondary recipients (supplemental Figure 7A). Intriguingly, IL1R1 KO demonstrated significantly longer survival and lower frequency of death irrespective of which of the LSK, CMP, or GMP populations was transplanted (supplemental Figure 7B). These data suggest that IL-1 signaling is crucial to LSPC activity in LSK/CMP/GMP populations in the MLL-AF9 model. In summary, loss of IL-1 signaling in the MLL-AF9 model of AML was associated with a more than five-fold decrease in LSC frequency, resulting from the restriction of activity to multiple LSPC populations from LSK to divergent CMP and GMP populations.

### UR241-2: A novel small molecule targeting interleukin-1 receptor-associated kinase 1/4 (IRAK1/4)

To inhibit aberrant IL-1 signaling in AML, we developed UR241-2, a novel small molecule inhibitor of IRAK1/4, guided by structure-activity relationship (SAR) (Figure 4A). IRAK1 and IRAK4 are serine-threonine kinases. In the canonical pathway, IRAK4 forms a complex with MyD88 and IRAK2 following IL1R1/TLR activation and induces downstream signaling through IRAK1 and TRAF6 (40) (Figure 2A). First, molecular docking studies using MedusaDock program (27, 28) showed superior affinity against IRAK1 (energy = -50.4 vs. -43.1 kcal/mol for UR241-2 vs. PF06650833); whereas for IRAK4, calculated affinities were much superior (energy = -51.3 vs. -39.9 kcal/mol for UR241-2 vs. PF06650833) (Figure 4B). Next, we performed kinase profiling assays, and UR241-2 showed a remarkable targeting ability against IRAK1 and IRAK4 (IC_50_ = 1.892e-09 M and 2.864e-09 M, respectively) (Figure 4C). Then, we conducted NanoBret assays using HEK293 cells, and UR241-2 demonstrated notably selective target engagement against IRAK4 (IC_50_ = 6.075e-09 M) than other targets (IC_50_ = >2.8e-07 M for others), including IRAK1 (IC_50_ = 6.621e-06 M) (Figure 4D). Lastly, in the kinome map analysis, UR241-2, at single dose (5 nM) screening, exhibited high target selectivity. Only 9 kinases in a panel of 682 kinases were inhibited by >25% activity, including IRAK1/4 (Figure 4E and supplemental Figure 8). These 9 kinases belong to STE (homologs of yeast Sterile 7, Sterile 11, and Sterile 20 kinases), Tyrosine Kinase (TK), and Tyrosine Kinase-Like (TKL) groups. Collectively, these data suggest that UR241-2 is a novel compound that selectively targets IRAK1/4, particularly IRAK4.

**Figure 4.**
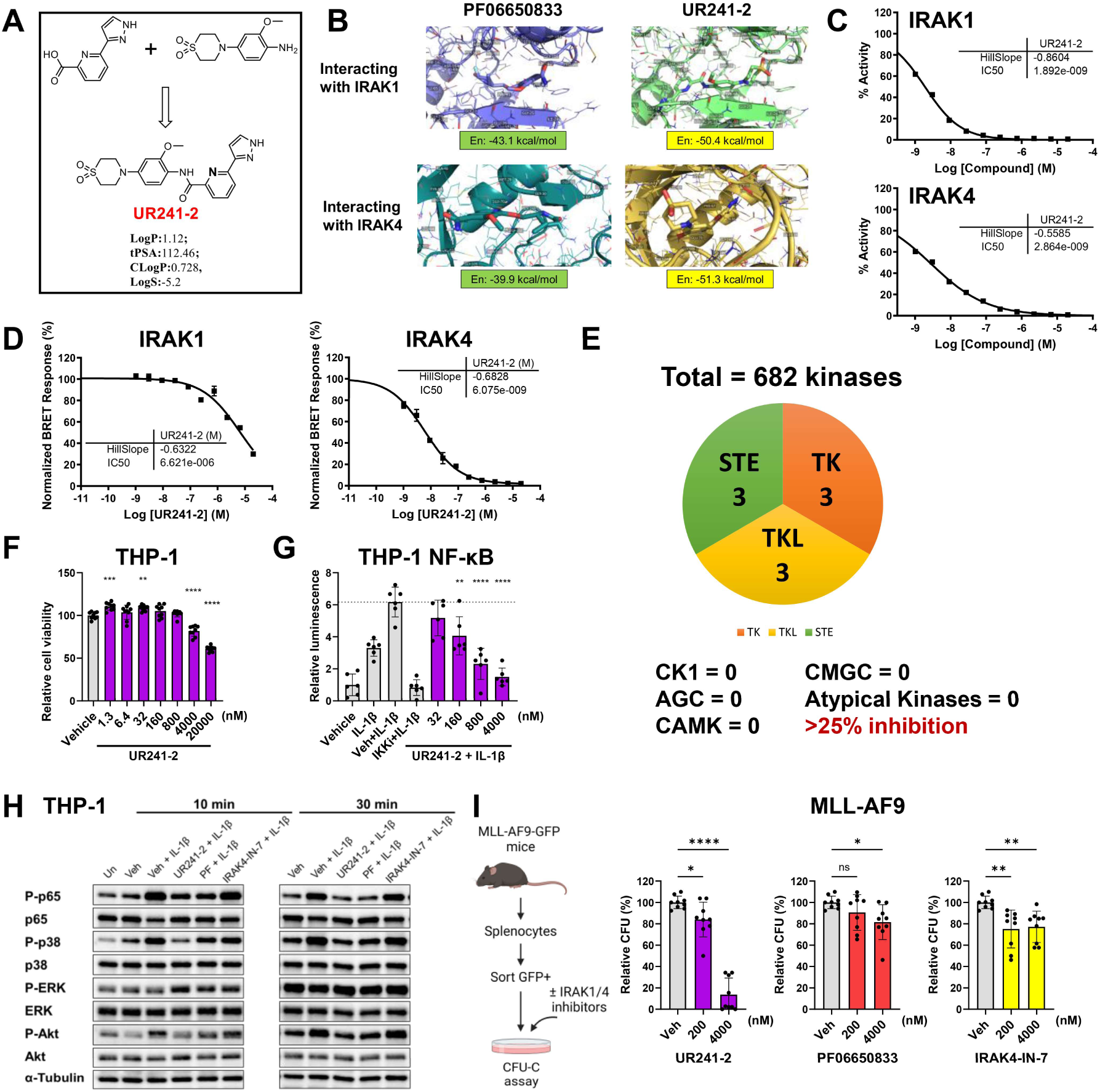
UR241-2: A novel small molecule targeting interleukin-1 receptor-associated kinase 1/4 (IRAK1/4). (A) Schematic of UR241-2 design and synthesis. This figure illustrates the logarithm of partition coefficient (LogP), topological polar surface area (tPSA), calculated logarithm of partition coefficient (CLogP), and logarithm of solubility (LogS). (B) Computer-simulated docking analyses for UR241-2 and PF06650833 against IRAK1 and IRAK4 using MedusaDock program. (C) The half maximal inhibitory concentration (IC_50_) of UR241-2 for IRAK1 and IRAK4 measured by kinase profiling assay with protein suspension. UR241-2 underwent testing in 10-dose IC_50_ duplicate mode, starting at 20 µM with 3-fold serial dilution. Reactions were carried out at 1 µM ATP. (D) The cellular IC_50_ of UR241-2 for IRAK1 and IRAK4 measured by NanoBRET Target Engagement Assay in HEK293 cells. HEK293 cells were transiently transfected with 1 µg IRAK1 and IRAK4 NanoLuc Fusion Vectors, along with 9 µg transfection carrier DNA. The transfected cells were treated with UR241-2, starting at 20 µM (10-dose with 3-fold dilution) for 1 hour. The target engagement was then quantified by the NanoBRET assay. (E) Pie chart depicting the kinome profiling of UR241-2. The chart highlights kinases with >25% inhibition by UR241-2 at a concentration of 5 nM. Only 9 kinases out of 682 kinases were affected. STE = Homologs of yeast Sterile 7, Sterile 11, and Sterile 20 kinases. TK = Tyrosine Kinase. TKL = Tyrosine Kinase-Like kinases. CK1 = Casein Kinase 1. AGC = PKA, PKG, and PKC families. CAMK = Calcium/calmodulin-dependent protein kinase. CMGC = CDK, MAPK, GSK3, and CLK families. (F) Cell viability assay of UR241-2 in THP-1 cells. Cells were treated with serial concentrations of UR241-2 (0.0013-20 μM) in the 96-well plate. After 48 hours, the cell viability was assessed using CellTiter-Glo 2.0 Cell Viability Assay. Vehicle = DMSO. (G) NF-κB reporter cell assay of UR241-2 in THP-1 NF-κB (Luc/GFP) reporter cells. Cells were pre-treated with serial concentrations of UR241-2 (0.032-4 μM) in the 96-well plate for 30 minutes and incubated with human IL-1β (10 ng/mL) for 6 hours. The activation levels of NF-κB were evaluated using ONE-Glo Luciferase Assay System. Vehicle = DMSO. IKKi = 20 μM IKK2 inhibitor (positive control). (H) Immunoblotting assay investigated IL-1 signaling in THP-1 cells treated with IRAK1/4 inhibitors. Cells were starved overnight in RPMI plus 0.5% FBS medium. Subsequently, cells were pre-treated with either vehicle (DMSO) or IRAK1/4 inhibitors (UR241-2, PF06650833, and IRAK4-IN-7; 4 μM) for 30 minutes and then incubated with human IL-1β (10 ng/mL) for 10 and 30 minutes. At each time point, cells were harvested and the phosphorylation and total levels of p65, p38, ERK, and Akt were analyzed. (I) CFU-C assays of MLL-AF9 cells treated with IRAK1/4 inhibitors (UR241-2, PF06650833, and IRAK4-IN-7). Live GFP^+^ MLL-AF9 splenocytes were sorted and mixed with drugs (0.2 and 4 μM) in the methylcellulose-based medium. Colonies were counted after 7 days of culture. Mean ± SD. Ordinary one-way ANOVA followed by Dunnett’s multiple comparisons test in *(F)*, *(G)*, and *(I)*. \**P* ≤ 0.05, \*\**P* ≤ 0.01, \*\*\**P* ≤ 0.001, \*\*\*\**P* ≤ 0.0001, ns = not significant. Schematics were created with BioRender.com.

### UR241-2 inhibits IL-1/IRAK1/4 signaling and colony-forming ability in AML

To functionally assess the impacts of UR241-2 in AML, we treated leukemia THP-1 cells with serial concentrations of UR241-2 and measured cell viability. UR241-2 does not induce acute cytotoxicity when treated with < 4 µM (resulting in ∼82% of viability after 48 hours of treatment) (Figure 4F). IL-1 is known to activate critical downstream effectors through IRAK1/4 such as NF-κB, p38, and ERK. To identify which downstream signaling mediators of IL-1/IRAK1/4 are impaired by UR241-2 in AML, we used leukemia THP-1 NF-κB reporter cells carrying luciferase. These cells were pre-incubated with UR241-2 for 30 minutes and stimulated with IL-1β for 6 hours. We demonstrated that UR241-2 can inhibit the activation of NF-κB following IL-1β stimulation in a dose-dependent manner (Figure 4G). We further used immunoblotting assays to validate the inhibition ability of UR241-2 in IL-1/IRAK1/4 signaling. PF06650833 and IRAK4-IN-7 are two IRAK4 inhibitors. IRAK4-IN-7 is an analog of CA-4948. CA-4948 has also been shown to be particularly effective in targeting the long isoform of IRAK4 (IRAK4-L) (16). We confirmed that UR241-2 robustly inhibits IL-1/IRAK1/4 signaling in AML cells including the phosphorylation of p65 and p38, following IL-1 stimulation (Figure 4H). Lastly, we investigated whether UR241-2 can suppress LSC function using colony-forming unit cell (CFU-C) assays of AML MLL-AF9 splenocytes treated with IRAK1/4 inhibitors. We found that UR241-2 significantly inhibits colony-forming ability in murine MLL-AF9 AML cells (Figure 4I), with similar or better efficacy compared to PF06650833 or IRAK4-IN-7. Taken together, our findings suggest that UR241-2 can potently inhibit the downstream signaling of IL-1/IRAK1/4 and suppress LSC function in AML.

### UR241-2 suppresses LSC function in primary human AML cells at diagnosis and relapse

We further assessed whether UR241-2 can abrogate LSC functions in primary human AML cells at diagnosis and relapse. No studies have compared the impact of IRAK1/4 inhibitors on the growth and survival of LSCs derived from paired diagnostic and relapsed AML patients. Patient AML25 sample has the t(9;11)(p22;q23) mutation (MLL-AF9 translocation). We demonstrated that UR241-2 suppresses LSC clonogenicity in primary human AML cells at diagnosis and relapse with similar or better efficacy compared to PF06650833 or IRAK4-IN-7 (Figure 5A-B and supplemental Figure 9A), as observed in the murine MLL-AF9 model (Figure 4I). Next, we determined that UR241-2, PF06650833, and IRAK4-IN-7 (0.2 and 4 μM) did not impair the colony-forming ability in NBM CD34^+^ cells (HSPC function) (Figure 5C). Finally, to investigate the impacts of UR241-2 on LSC engraft ability *in vivo*, we treated murine MLL-AF9 GFP^+^ splenocytes with vehicle, UR241-2, and PF06650833 *ex vivo* for three days and transplanted these cells into recipient mice (Figure 5D). After four weeks, we observed that UR241-2 significantly suppresses the engraftment levels of AML cells (live GFP^+^ cells) in BM (Figure 5E) and PB (supplemental Figure 9B). It also decreases the WBC in PB and increases the RBC, HGB, and PLT counts compared to the vehicle control group (supplemental Figure 9C). AML MLL-AF9 mice treated with vehicle or PF06650833 have splenomegaly; however, the spleen weights were significantly decreased in the UR241-2 group at the time of sacrifice (Figure 5F). In summary, our findings demonstrate that targeting IRAK1/4 using the novel compound UR241-2 is able to impair LSC clonogenicity and engraftment capability in AML following treatment and disease progression. This provides a potential therapeutic strategy to eradicate the LSC population at both diagnosis and relapse and improve the outcomes of AML patients.

**Figure 5.**
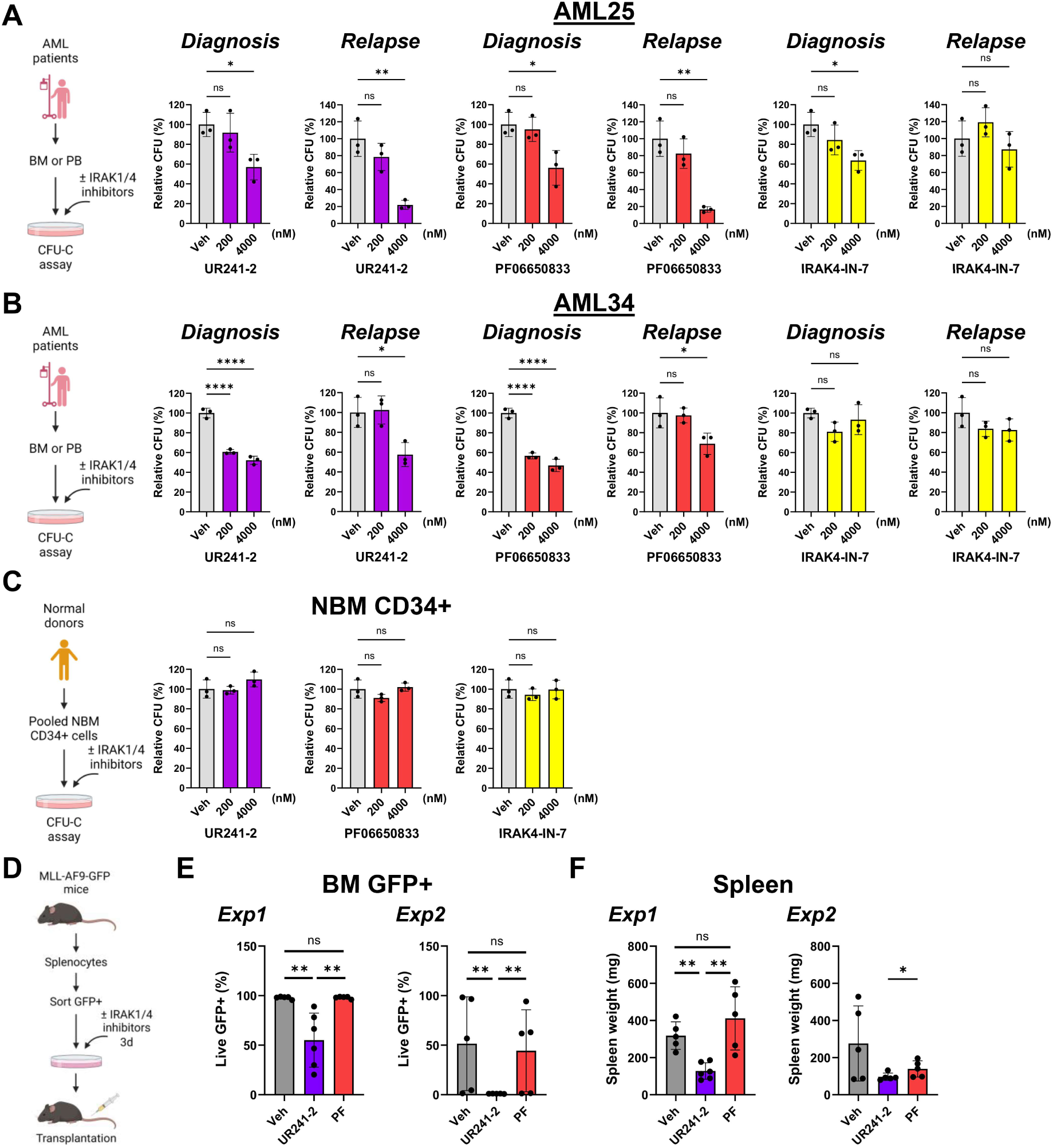
UR241-2 suppresses LSC function in primary human AML cells at diagnosis and in relapse. (A) CFU-C assays of paired primary human AML cells at diagnosis and in relapse (AML25). Cells were treated with IRAK1/4 inhibitors UR241-2, PF06650833, and IRAK4-IN-7 (0.2 and 4 μM) in methylcellulose-based medium. (B) CFU-C assays of paired diagnostic and relapsed AML34 cells, as described in *(A).* (C) CFU-C assays of primary human NBM CD34^+^ cells treated with UR241-2, PF06650833, and IRAK4-IN-7 (0.2 and 4 μM) in methylcellulose-based medium. Mean ± SD. Ordinary one-way ANOVA followed by Dunnett’s multiple comparisons test in *(A-C)*. (D) Schematic of UR241-2 *ex vivo* treatment and transplantation using MLL-AF9 AML model. MLL-AF9 GFP^+^ splenocytes were sorted and treated with vehicle (DMSO), UR241-2, and PF06650833 (4 μM) *ex vivo* for three days. These cells were transplanted into recipient C57BL/6J mice following treatment. Mice were sacrificed and assessed after four weeks. (E) Engraftment levels of MLL-AF9 GFP^+^ cells in the bone marrow in recipients at four weeks after transplantation as described in *(D)*. Mean ± SD. Two-tailed Mann-Whitney test. (F) Spleen weights in recipients at four weeks after transplantation as described in *(D)*. Mean ± SD. Two-tailed Mann-Whitney test. \**P* ≤ 0.05, \*\**P* ≤ 0.01, \*\*\*\**P* ≤ 0.0001, ns = not significant. Schematics were created with BioRender.com.

## Discussion

Pre-leukemic and LSC populations are often associated with relapse after initial treatment for AML (42-44). Our previous research has elucidated the evolution of LSC populations during AML treatment and disease progression, revealing large-scale changes in LSC frequency, phenotype, and genetic stability (5). These findings underscore the critical link between LSC clones and leukemogenesis. Therefore, it is imperative to intervene with therapies that eradicate LSCs early in the disease course and throughout all stages. Nevertheless, the underlying mechanisms driving this phenomenon are not fully understood. In this study, we employed TLDA transcriptome analysis and immunophenotyping to identify the upregulation of IL-1 signaling in LSCs among AML patients during treatment and subsequent relapse. By utilizing CFU-C and xenotransplantation assays, we demonstrated that loss of IL-1 signaling components, such as IL1RAP and IL1R1, profoundly impacts LSC colony-forming ability and engraftment capability. These findings underscore the significance of the leukemia microenvironment and its inflammatory milieu on disease initiation and progression.

Pro-inflammatory cytokines, including IL-1, IL-6, or TNF-α, are elevated within the bone marrow microenvironment (BMME) in MDS and AML (45, 46). A proposed model suggests that MDS progression within the inflammatory BMME begins with a mutation in HSPCs, altering their response to inflammation. Over time, pre-leukemic and MDS HSPCs outcompete normal HSPCs, accumulate more mutations, and eventually dominate the blood system, leading to impaired hematopoiesis at the MDS stage (10, 47). Furthermore, IL-1 signaling mediates alterations in the abundance and function of BMME cell populations, including mesenchymal stromal cells, and drives hematopoiesis skewing during aging (48, 49). A transcriptomic analysis has revealed an association between pro-inflammatory mediators, including *IL1R1*, and short event-free survival of AML patients (50). In our study, we have unveiled the critical involvement of IL-1 signaling in AML patients experiencing relapse following conventional intensive “7+3” therapy. It would be intriguing to further investigate whether IL-1 signaling similarly plays a pivotal role in AML patients undergoing other novel therapies, such as azacitidine and venetoclax, to drive relapse.

Recent studies and clinical trials have focused on IRAK1 and IRAK 4 in AML and MDS, leading to the development of new drugs targeting these pathways (16, 17, 23, 51-54). IRAK1 has been shown to be overexpressed and activated in MDS BM samples. Genetic and pharmacological inhibition of IRAK1 effectively suppresses MDS cells (11, 51). IRAK4 exists in two isoforms, the short isoform (IRAK4-S) and the long isoform (IRAK-L). Common spliceosome mutations found in AML and MDS, such as *U2AF1* or *SF3B1*, can promote alternate splicing to include exon 4 or exon 6, respectively, resulting in increased expression of IRAK4-L. This upregulation leads to significant NF-kB activation, supporting the growth of AML and MDS cells. CA-4948 has demonstrated specific efficacy in targeting IRAK4-L, effectively inhibiting leukemia cell growth in both AML and MDS (11, 16, 17).

While current clinical trials of IRAK4 inhibitors have shown promise in treating AML and MDS, the overall response remains limited (23, 55). A recent study has highlighted the insufficiency of relying solely on single IRAK4 treatment due to functional complementation and compensation by its paralog IRAK1 (54). This suggests that co-targeting IRAK1 and IRAK4 may achieve better efficiency in eliminating LSPCs (54). Our findings highlight the potency of UR241-2, a novel inhibitor that simultaneously targets IRAK1 and IRAK4. This compound effectively disrupts the function of LSPCs in both diagnostic and relapsed AML patient samples. Future studies are warranted to thoroughly explore the effects of UR241-2 on AML LSCs and MDS HSPCs at various disease stages. Comprehensive analyses of transcriptome and proteome signatures are needed to identify potential resistance mechanism against IRAK1/4 inhibitors including UR241-2. These investigations hold promise for optimizing current therapeutic strategies in AML and MDS.

## Acknowledgments

The authors thank the University of Rochester Medical Center (URMC) Flow Cytometry Shared Resource for maintaining fluorescence-activated cell sorting and flow cytometry facilities; the UR Genomics Research Center for assisting with transcriptome analyses; and Naxin Guo (URMC), Yuko Kawano (URMC), and Rita Assi (IUSM) for their technical support, valuable insights, and suggestions. The authors also thank Jessica Mavor and Laurie Ford for their support in obtaining patient samples at the Wilmot Cancer Institute and Roswell Park Comprehensive Cancer Center. We express our gratitude to the patients and their families who participated in the studies from which all samples were obtained.

This study was supported in part by the National Institutes of Health National Cancer Institute (R21CA149848 and R01CA166280), Leukemia & Lymphoma Society (7020-19), J. P. Wilmot Cancer Center Pilot Award, University of Rochester Technologies Development Fund, University of Rochester CTSI, and Empire Discovery Institute. N.V.D acknowledges the support from the National Institutes of Health (R35GM134864) and the Passan Foundation. This project was supported by the Penn State College of Medicine’s Artificial Intelligence and Biomedical Informatics Program.

## Authorship

### Contribution

T.-C.H. and M.W.B. designed the study; T.-C.H., M.W.L., H.K., D.K.B., E.A.L., and R.K.S. performed experiments; T.-C.H., M.W.L., H.K., Y.-C.C., C.C., J.W., N.V.D., and R.K.S. performed analyses; L.M.C., J.L.L., and C.T.J. contributed vital new reagents or analytical tools; T.-C.H. wrote the manuscript; T.-C.H., J.L.L., C.T.J., R.K., and M.W.B. revised the manuscript; all authors approved the final version of the manuscript.

### Conflict-of-interest Statement

M.W.B., R.K.S., and L.M.C. hold the patent for UR241-2 (CA3186896A1). The remaining authors have declared that no conflict of interest exists.

## Notes

### Competing Interest Statement

The authors have declared no competing interest.

## References

1. Jaramillo S, and Schlenk RF. Update on current treatments for adult acute myeloid leukemia: to treat acute myeloid leukemia intensively or non-intensively? That is the question. Haematologica. 2023;108(2):342–52.

2. Dohner H, Weisdorf DJ, and Bloomfield CD. Acute Myeloid Leukemia. N Engl J Med. 2015;373(12):1136–52.

3. Kantarjian H, Kadia T, DiNardo C, Daver N, Borthakur G, Jabbour E, et al. Acute myeloid leukemia: current progress and future directions. Blood Cancer J. 2021;11(2):41.

4. Stauber J, Greally JM, and Steidl U. Preleukemic and leukemic evolution at the stem cell level. Blood. 2021;137(8):1013–8.

5. Ho TC, LaMere M, Stevens BM, Ashton JM, Myers JR, O’Dwyer KM, et al. Evolution of acute myelogenous leukemia stem cell properties after treatment and progression. Blood. 2016;128(13):1671–8.

6. Dohner H, Wei AH, Appelbaum FR, Craddock C, DiNardo CD, Dombret H, et al. Diagnosis and management of AML in adults: 2022 recommendations from an international expert panel on behalf of the ELN. Blood. 2022;140(12):1345–77.

7. Dohner H, Wei AH, and Lowenberg B. Towards precision medicine for AML. Nat Rev Clin Oncol. 2021;18(9):577–90.

8. Daver N, Wei AH, Pollyea DA, Fathi AT, Vyas P, and DiNardo CD. New directions for emerging therapies in acute myeloid leukemia: the next chapter. Blood Cancer J. 2020;10(10):107.

9. Thol F, and Ganser A. Treatment of Relapsed Acute Myeloid Leukemia. Curr Treat Options Oncol. 2020;21(8):66.

10. Trowbridge JJ, and Starczynowski DT. Innate immune pathways and inflammation in hematopoietic aging, clonal hematopoiesis, and MDS. J Exp Med. 2021;218(7).

11. Vegivinti CTR, Keesari PR, Veeraballi S, Martins Maia CMP, Mehta AK, Lavu RR, et al. Role of innate immunological/inflammatory pathways in myelodysplastic syndromes and AML: a narrative review. Exp Hematol Oncol. 2023;12(1):60.

12. Sims JE, and Smith DE. The IL-1 family: regulators of immunity. Nat Rev Immunol. 2010;10(2):89–102.

13. Fitzgerald KA, and Kagan JC. Toll-like Receptors and the Control of Immunity. Cell. 2020;180(6):1044–66.

14. Weber A, Wasiliew P, and Kracht M. Interleukin-1 (IL-1) pathway. Sci Signal. 2010;3(105):cm1.

15. Pietras EM, Mirantes-Barbeito C, Fong S, Loeffler D, Kovtonyuk LV, Zhang S, et al. Chronic interleukin-1 exposure drives haematopoietic stem cells towards precocious myeloid differentiation at the expense of self-renewal. Nat Cell Biol. 2016;18(6):607–18.

16. Smith MA, Choudhary GS, Pellagatti A, Choi K, Bolanos LC, Bhagat TD, et al. U2AF1 mutations induce oncogenic IRAK4 isoforms and activate innate immune pathways in myeloid malignancies. Nat Cell Biol. 2019;21(5):640–50.

17. Choudhary GS, Pellagatti A, Agianian B, Smith MA, Bhagat TD, Gordon-Mitchell S, et al. Activation of targetable inflammatory immune signaling is seen in myelodysplastic syndromes with SF3B1 mutations. Elife. 2022;11.

18. Sallman DA, and List A. The central role of inflammatory signaling in the pathogenesis of myelodysplastic syndromes. Blood. 2019;133(10):1039–48.

19. Barreyro L, Will B, Bartholdy B, Zhou L, Todorova TI, Stanley RF, et al. Overexpression of IL-1 receptor accessory protein in stem and progenitor cells and outcome correlation in AML and MDS. Blood. 2012;120(6):1290–8.

20. Carey A, Edwards DKt, Eide CA, Newell L, Traer E, Medeiros BC, et al. Identification of Interleukin-1 by Functional Screening as a Key Mediator of Cellular Expansion and Disease Progression in Acute Myeloid Leukemia. Cell Rep. 2017;18(13):3204–18.

21. Broderick L, and Hoffman HM. IL-1 and autoinflammatory disease: biology, pathogenesis and therapeutic targeting. Nat Rev Rheumatol. 2022;18(8):448–63.

22. Dinarello CA. Overview of the IL-1 family in innate inflammation and acquired immunity. Immunol Rev. 2018;281(1):8–27.

23. Bennett J, and Starczynowski DT. IRAK1 and IRAK4 as emerging therapeutic targets in hematologic malignancies. Curr Opin Hematol. 2022;29(1):8–19.

24. Majeti R, Becker MW, Tian Q, Lee TL, Yan X, Liu R, et al. Dysregulated gene expression networks in human acute myelogenous leukemia stem cells. Proc Natl Acad Sci U S A. 2009;106(9):3396–401.

25. Eppert K, Takenaka K, Lechman ER, Waldron L, Nilsson B, van Galen P, et al. Stem cell gene expression programs influence clinical outcome in human leukemia. Nat Med. 2011;17(9):1086–93.

26. Ashton JM, Balys M, Neering SJ, Hassane DC, Cowley G, Root DE, et al. Gene sets identified with oncogene cooperativity analysis regulate in vivo growth and survival of leukemia stem cells. Cell Stem Cell. 2012;11(3):359–72.

27. Wang J, and Dokholyan NV. MedusaDock 2.0: Efficient and Accurate Protein-Ligand Docking With Constraints. J Chem Inf Model. 2019;59(6):2509–15.

28. Ding F, Yin S, and Dokholyan NV. Rapid flexible docking using a stochastic rotamer library of ligands. J Chem Inf Model. 2010;50(9):1623–32.

29. Yin S, Biedermannova L, Vondrasek J, and Dokholyan NV. MedusaScore: an accurate force field-based scoring function for virtual drug screening. J Chem Inf Model. 2008;48(8):1656–62.

30. Metz KS, Deoudes EM, Berginski ME, Jimenez-Ruiz I, Aksoy BA, Hammerbacher J, et al. Coral: Clear and Customizable Visualization of Human Kinome Data. Cell Syst. 2018;7(3):347–50 e1.

31. Saito Y, Kitamura H, Hijikata A, Tomizawa-Murasawa M, Tanaka S, Takagi S, et al. Identification of therapeutic targets for quiescent, chemotherapy-resistant human leukemia stem cells. Sci Transl Med. 2010;2(17):17ra9.

32. Shlush LI, Mitchell A, Heisler L, Abelson S, Ng SWK, Trotman-Grant A, et al. Tracing the origins of relapse in acute myeloid leukaemia to stem cells. Nature. 2017;547(7661):104-8.

33. Subramanian A, Tamayo P, Mootha VK, Mukherjee S, Ebert BL, Gillette MA, et al. Gene set enrichment analysis: a knowledge-based approach for interpreting genome-wide expression profiles. Proc Natl Acad Sci U S A. 2005;102(43):15545–50.

34. Ng SW, Mitchell A, Kennedy JA, Chen WC, McLeod J, Ibrahimova N, et al. A 17-gene stemness score for rapid determination of risk in acute leukaemia. Nature. 2016;540(7633):433-7.

35. Mitchell K, Barreyro L, Todorova TI, Taylor SJ, Antony-Debre I, Narayanagari SR, et al. IL1RAP potentiates multiple oncogenic signaling pathways in AML. J Exp Med. 2018;215(6):1709–27.

36. Dinarello CA, Simon A, and van der Meer JW. Treating inflammation by blocking interleukin-1 in a broad spectrum of diseases. Nat Rev Drug Discov. 2012;11(8):633–52.

37. Askmyr M, Agerstam H, Hansen N, Gordon S, Arvanitakis A, Rissler M, et al. Selective killing of candidate AML stem cells by antibody targeting of IL1RAP. Blood. 2013;121(18):3709–13.

38. Jaras M, Johnels P, Hansen N, Agerstam H, Tsapogas P, Rissler M, et al. Isolation and killing of candidate chronic myeloid leukemia stem cells by antibody targeting of IL-1 receptor accessory protein. Proc Natl Acad Sci U S A. 2010;107(37):16280–5.

39. Agerstam H, Hansen N, von Palffy S, Sanden C, Reckzeh K, Karlsson C, et al. IL1RAP antibodies block IL-1-induced expansion of candidate CML stem cells and mediate cell killing in xenograft models. Blood. 2016;128(23):2683–93.

40. Rhyasen GW, and Starczynowski DT. IRAK signalling in cancer. Br J Cancer. 2015;112(2):232–7.

41. Krivtsov AV, Twomey D, Feng Z, Stubbs MC, Wang Y, Faber J, et al. Transformation from committed progenitor to leukaemia stem cell initiated by MLL-AF9. Nature. 2006;442(7104):818-22.

42. Pollyea DA, and Jordan CT. Therapeutic targeting of acute myeloid leukemia stem cells. Blood. 2017;129(12):1627–35.

43. Shastri A, Will B, Steidl U, and Verma A. Stem and progenitor cell alterations in myelodysplastic syndromes. Blood. 2017;129(12):1586–94.

44. Ishikawa F, Yoshida S, Saito Y, Hijikata A, Kitamura H, Tanaka S, et al. Chemotherapy-resistant human AML stem cells home to and engraft within the bone-marrow endosteal region. Nat Biotechnol. 2007;25(11):1315–21.

45. Banerjee T, Calvi LM, Becker MW, and Liesveld JL. Flaming and fanning: The Spectrum of inflammatory influences in myelodysplastic syndromes. Blood Rev. 2019;36:57–69.

46. Luciano M, Krenn PW, and Horejs-Hoeck J. The cytokine network in acute myeloid leukemia. Front Immunol. 2022;13:1000996.

47. Pietras EM. Inflammation: a key regulator of hematopoietic stem cell fate in health and disease. Blood. 2017;130(15):1693–8.

48. Frisch BJ, Hoffman CM, Latchney SE, LaMere MW, Myers J, Ashton J, et al. Aged marrow macrophages expand platelet-biased hematopoietic stem cells via Interleukin1B. JCI Insight. 2019;5.

49. Mitchell CA, Verovskaya EV, Calero-Nieto FJ, Olson OC, Swann JW, Wang X, et al. Stromal niche inflammation mediated by IL-1 signalling is a targetable driver of haematopoietic ageing. Nat Cell Biol. 2023;25(1):30–41.

50. Stratmann S, Yones SA, Garbulowski M, Sun J, Skaftason A, Mayrhofer M, et al. Transcriptomic analysis reveals proinflammatory signatures associated with acute myeloid leukemia progression. Blood Adv. 2022;6(1):152–64.

51. Rhyasen GW, Bolanos L, Fang J, Jerez A, Wunderlich M, Rigolino C, et al. Targeting IRAK1 as a therapeutic approach for myelodysplastic syndrome. Cancer Cell. 2013;24(1):90–104.

52. Melgar K, Walker MM, Jones LM, Bolanos LC, Hueneman K, Wunderlich M, et al. Overcoming adaptive therapy resistance in AML by targeting immune response pathways. Sci Transl Med. 2019;11(508).

53. Muto T, Walker CS, Agarwal P, Vick E, Sampson A, Choi K, et al. Inactivation of p53 provides a competitive advantage to del(5q) myelodysplastic syndrome hematopoietic stem cells during inflammation. Haematologica. 2023;108(10):2715–29.

54. Bennett J, Ishikawa C, Agarwal P, Yeung J, Sampson A, Uible E, et al. Paralog-specific signaling by IRAK1/4 maintains MyD88-independent functions in MDS/AML. Blood. 2023;142(11):989–1007.

55. Garcia-Manero G, Winer ES, DeAngelo DJ, Tarantolo S, Sallman DA, Dugan J, et al. S129: TAKEAIM LEUKEMIA-A PHASE 1/2A STUDY OF THE IRAK4 INHIBITOR EMAVUSERTIB (CA-4948) AS MONOTHERAPY OR IN COMBINATION WITH AZACITIDINE OR VENETOCLAX IN RELAPSED/REFRACTORY AML OR MDS. HemaSphere. 2022;6:30–1.

