## Supplemental Data for "Targeting IL-1/IRAK1/4 signaling in Acute Myeloid Leukemia Stem Cells Following Treatment and Relapse"

**This Supplemental File includes:**

Supplemental Methods
Supplemental Figures 1 to 9
Supplemental Tables 1 to 3
Supplemental References

### Supplemental Methods

#### Primary human samples

Primary human samples from AML patients or normal donors were obtained following informed consent under institutional review board approved protocols and the methods outlined in our previous work (1). In brief, for AML patient samples, any subject with primary or secondary AML who is receiving cytotoxic therapy and/or allogeneic bone marrow/peripheral blood (BM/PB) stem cell transplant is eligible. For normal donor samples, any individual over the age of 18 and in good health is eligible. BM aspirates, apheresis product, or PB were collected from the University of Rochester Medical Center (URMC) and Roswell Park Comprehensive Cancer Center. Mononuclear cells (MNCs) were isolated using Ficoll-Paque Plus (MilliporeSigma) and stored in liquid nitrogen. AML patient characteristics are detailed in supplemental Table 1. Primary human cells were cultured in serum free medium (SFM) supplemented with cytokine mixture (cytokine SFM). SFM contains IMDM (Gibco), 20% BIT-9500 (STEMCELL Technologies), 1x low density lipoprotein (LDL, Calbiochem), 0.05 mM 2-mercaptoethanol (Gibco), 1x L-Glutamine (Gibco), and 1x penicillin-streptomycin (Pen-Strep) (Gibco). The cytokine mixture contains recombinant human (rh) SCF (25 ng/mL), rh IL-3 (10 ng/mL), rh IL-7 (10 ng/mL), rh FLT3-Ligand (10 ng/mL), and rh GCSF (10 ng/mL) (PeproTech).

#### Cell lines

Human leukemic cell lines THP-1 (ATCC TIB-202; RRID:CVCL_0006), U937 (ATCC CRL-1593.2; RRID:CVCL_0007), and KG-1a (ATCC CCL-246.1; RRID:CVCL_1824) were obtained from ATCC. THP-1 cells were cultured in RPMI 1640 (Gibco) supplemented with 10% heat-inactivated fetal bovine serum (FBS) (Corning), 0.05 mM 2-mercaptoethanol, and 1x Pen-Strep. U937 cells were cultured in RPMI 1640 supplemented with 10% FBS and 1x Pen-Strep. KG-1a cells were cultured in IMDM (Gibco) supplemented with 10% FBS and 1x Pen-Strep. 293TN cells (RRID:CVCL_UL49) were obtained from System Biosciences and cultured in DMEM (Gibco) supplemented with 10% FBS and 1x Glutamax (Gibco) (1x Pen-Strep was added if antibiotics were required). All cells were cultured at 37°C and 5% $CO_2$. Cell lines were directly purchased from ATCC or System Biosciences, divided into aliquots, and used at early passages.

#### Animals

C57BL/6J (B6; RRID:IMSR_JAX:000664), B6.SJL-*Ptprc*$^{a}$ *Pepc*$^{b}$/BoyJ (Pep Boy), B6.129S7-*Il1r1*$^{tm1Imx}$/J (Il1r1$^{-}$; IL1R1 KO), and NOD.Cg-*Prkdc*$^{scid}$ *Il2rg*$^{tm1Wjl}$/SzJ (NSG) mice were purchased from Jackson lab or bred in-house at URMC. Animal experiments in this study were approved by the Institutional Animal Care and Use Committee (IACUC) at the URMC.

**Flow cytometry analysis**

To perform immunophenotyping, primary AML or NBM cells were incubated with antibodies against human CD34 (or CD32), CD38, and IL1RAP (or IL1R1) in FACS buffer (1x PBS plus 0.5% FBS or 2% FBS) at 4 °C for 20-30 minutes. Antibodies used in the study are detailed in supplemental Table 3. Cells were then washed and stained with 1x 4', 6 diamidino-2-phenylindole (DAPI) (Invitrogen) in FACS buffer. Flow cytometry analyses were conducted using a LSR II or LSRFortessa system (BD Biosciences) and data analyses were conducted using FlowJo software (BD; RRID:SCR_008520).

**Fluorescence-activated cell sorting (FACS)**

To prepare samples for TaqMan Low Density Array based analyses, primary human AML cells were stained with antibodies against human CD3, CD34 (or CD32), and CD38. Normal HSCs were isolated from human NBM samples enriched for CD34$^{+}$ cells using the MACS Indirect CD34 MicroBead Kit (Miltenyi Biotec). Enriched CD34$^{+}$ cells were stained with biotinylated lineage antibodies (against human CD3, CD14, CD15, CD16, CD19, and CD56), as well as streptavidin, antibodies against human CD34, CD38, CD90, CD123, glycophorin A, and 1x DAPI. Live AML LSCs (CD3$^{-}$CD34/CD38 or CD3$^{-}$CD32/CD38) and NBM HSCs (CD34$^{+}$CD38$^{-}$CD90$^{+}$Lin$^{-}$) were double-sorted into TRIzol Reagent (Invitrogen) using a BD FACSAria cell sorter (BD Biosciences).

**Real-Time Quantitative PCR (RT-qPCR)**

Total RNA was extracted using TRIzol Reagent, RNeasy Mini Kit, RNeasy Micro Kit, RNeasy Plus Mini Kit, or RNeasy Plus Micro Kit (Qiagen). cDNA was synthesized with iScript cDNA synthesis Kit. For individual gene analyses, RT-qPCR was performed using TaqMan Gene Expression Assay, TaqMan PreAmp Master Mix, and TaqMan Gene Expression Master Mix (Thermo Fisher Scientific). The reactions were carried out using the LightCycler 480 Real-Time

PCR System (Roche) or the Applied Biosystems QuantStudio 12K Flex Real-Time PCR System (Thermo Fisher Scientific). The relative quantification analyses were performed using *GAPDH* and *HPRT1* as reference controls.

**Lentiviral production**

The lentiviral cloning, production, and titer determination was performed as previously described (2). In brief, short hairpin RNA (shRNA) oligonucleotides targeting IL1RAP and IL1R1 as well as scramble shRNA (sh-IL1RAP: GCTAGTGGTGATTCTCATTGT; sh-IL1R1: GCAAATAGCCATGTATAATGC; sh-SCR: scramble shRNA as control) were constructed into pLKO.1-GFP vector. The lentivector pLKO.1-GFP construct DNA, pPax2 (packaging plasmid), and pMD2.G (vesicular stomatitis virus-g envelop plasmid) were transfected into 293TN cells using TransIT-LT1 Transfection Reagent (Mirus Bio) following the instructions from the manufacturer. The transfection medium was replaced with fresh 293TN medium (DMEM + 10% FBS + 1% Glutamax) on the following day. Lentiviral particles in the supernatant were harvested on the next two days and filtered through 0.45 µm PES syringe filter (Nalgene). Pooled lentiviral supernatant was either directly aliquoted in cryovials (Corning) or PEG precipitated before being aliquoted, and flash-frozen using liquid nitrogen, followed by storage at -80°C. Lentiviral titer was determined, and high-titer lentiviral particles were produced for subsequent assays.

**Retroviral production**

The retroviral production and titer determination were carried out as previously described (2-4). Briefly, the mouse stem cell virus (MSCV) retroviral plasmid containing MLL-AF9-GFP oncogene (pMSCV-MLL-AF9-GFP) was transfected into Platinum-E cells (a retroviral packaging cell line; Cell Biolabs), using either the calcium phosphate solution with calcium chloride and HEPES buffered saline or the TransIT-LT1 Transfection Reagent (Mirus Bio). After two days of transfection, retroviral particles were harvested, filtered, and then flash frozen using liquid nitrogen, followed by storage at -80°C.

**Infection of human primary cells and leukemia cell lines**

To infect primary human cells, thawed primary AML cells were cultured in cytokine SFM for 5 to 6 hours before infection. For NBM CD34$^{+}$ cells, three NBM samples were equally pooled and enriched using the MACS Indirect CD34 MicroBead Kit. These $CD34^+$ cells were then cultured in cytokine SFM for 5 to 6 hours before infection. A total of $5 \times 10^6$ primary AML or enriched normal $CD34^+$ cells were resuspended in 500 μL cytokine SFM and mixed with 50 μL PEG-concentrated viral supernatant along with 5 μL polybrene (AmericanBio). To infect human leukemia cell lines, 400 μL of non-concentrated viral supernatant and 8 μL polybrene was added into $1 \times 10^6$ - $2 \times 10^6$ cells in 400 μL complete medium. The infection efficiency was evaluated using flow cytometry on an LSR II or LSRFortessa system (BD Biosciences). Live $GFP^+$ AML cells ($DAPI^-GFP^+$ or $DAPI^-GFP^+CD3^-$), NBM $CD34^+$ cells ($DAPI^-GFP^+CD34^+$), and $GFP^+$ leukemia cells ($DAPI^-GFP^+$) were sorted under sterile conditions at 2 to 3 days (typically 3 days) post infection using a BD FACSAria cell sorter (BD Biosciences) for following assays.

**Generation of murine MLL-AF9 AML model**

The murine MLL-AF9 AML model was generated as previously described (4, 5). In brief, $Lin^-Sca\text{-}1^+c\text{-}Kit^+$ (LSK) BM cells (Lin = CD3e, B220, GR-1, and Ter119) from WT and IL1R1 KO mice (C57BL/6J background) were sorted and cultured in LSK medium (alpha-MEM supplemented with 10% FBS and 50 ng/mL recombinant murine SCF, Flt3-Ligand, and IL-6 (all cytokines sourced from PeproTech)). Next, cells were infected with retroviral supernatant, containing the leukemic MLL-AF9-GFP vector, twice per day for three days. The infected cells were then transplanted into lethally irradiated Pep Boy recipient mice (5 Gy × 2, twenty-four hours apart) through tail-vein injection. Throughout the study, mice were monitored routinely for their complete blood count (CBC) values and engraftment levels (live $GFP^+$ cells) in peripheral blood (PB). When the mice appeared moribund, they were sacrificed, and engraftment levels in BM, spleen, and PB were measured using flow cytometry (GFP, CD45.1, or CD45.2). BM and spleen cells from leukemic mice were harvested for subsequent assays.

**LSK/CMP/GMP transplantation assay**

Splenocytes from primary transplantations of MLL-AF9 IL1R1 WT and MLL-AF9 IL1R1 KO mice (three mice per group) were pooled and sorted for leukemia LSK, CMP, and GMP populations (LSK: $Lin^-Sca\text{-}1^+c\text{-}Kit^+$; CMP: $Lin^-c\text{-}Kit^+(16/32)FC\gamma R^-CD34^+$; GMP: $Lin^-c\text{-}Kit^+(16/32)FC\gamma R^+CD34^+$) (Lin = CD3e, B220, GR-1, and Ter119). The number of cells for each population was determined based on the proportion of cells identified during sorting, with the smallest population serving as the reference. Additionally, fresh CD45.1 BM competitor cells were mixed with the test cells to achieve a total of $1 \times 10^6$ cells per mouse before transplantation. Finally, the cells were injected into lethally irradiated Pep Boy recipient mice (5 Gy × 2) via the tail-vein route.

**Colony-forming unit cell (CFU-C) assay**

CFU-C assays were conducted using different cells. For human cells, including primary AML, NBM CD34$^+$ cells, and leukemia cell lines, cells were plated in MethoCult GF H4534 (STEMCELL Technologies) supplemented with human EPO (3 U/mL), GCSF (10 ng/mL), and 1x Pen-Strep on a 35 mm petri dish (Corning) at appropriate cell densities ($1 \times 10^4$ cells/mL for primary AML cells, 2.5-5 × $10^3$ cells/mL for NBM CD34$^+$ cells, and 1-10 × $10^3$ cells/mL for U937 and THP-1 cells). Colonies were scored after 7-21 days of culture (typically 14 days). For murine cells, live GFP$^+$ MLL-AF9 splenocytes (DAPI$^-$GFP$^+$) were sorted and plated in MethoCult GF M3534 (STEMCELL Technologies) plus 1x Pen-Strep on a 35 mm petri dish ($5 \times 10^3$ cells/mL). Colonies were scored after 7 days of culture.

**Cell viability assay**

THP-1 cells were seeded in 96-well plates ($1 \times 10^5$ cells/well) in complete medium and treated with UR241-2 at the indicated concentrations. After incubating for 24 and 48 hours, cell viability was assessed using the CellTiter-Glo Luminescent Cell Viability Assay (Promega) following the manufacturer’s instructions.

**NF-κB reporter cell assay**

To generate mCherry NF-κB (GFP/Luc) reporter cells, we followed the lentiviral protocols described above and in a previous report (2). In brief, we constructed pGreenFire Lenti-Reporter genes (SBI) into the pLKO.1-mCherry vector to create the mCherry NF-κB (GFP/Luc) reporter lentivectors. High-titer lentiviral particles were produced and used to infect AML THP-1 cells. After 3 days of incubation, live mCherry$^+$ cells were sorted to obtain the THP-1 NF-κB reporter cells. For drug treatment, THP-1 NF-κB reporter cells were seeded in 96-well plates ($1 \times 10^5$ cells/well) in RPMI medium plus 0.5% FBS and pre-treated with drugs at the indicated concentrations for 30 minutes (UR241-2: 0.032-4 μM; IKK2 inhibitor (IKKi, positive control):

20 μM). Subsequently, human IL-1β (10 ng/mL) was added to the plates and incubated for another 6 hours. The activation of NF-κB in cells was assessed using the ONE-Glo Luciferase Assay System (Promega).

**Immunoblotting**

Protein lysates were prepared by lysing cells in 1x Cell Lysis Buffer (Cell Signaling Technology) supplemented with 1x Protease/Phosphatase Inhibitor Cocktail (Cell Signaling Technology). Protein concentration was quantified using the Pierce BCA Protein Assay Kit (Thermo Scientific). Samples were mixed with 2x or 4x Laemmli Sample Buffer (Bio-Rad) and 5% 2-mercaptoethanol (Sigma-Aldrich) before undergoing protein electrophoresis and blotting according to the manufacturer's instructions (Bio-Rad). Alternatively, cells were lysed in 1x SDS sample buffer for subsequent assays. Any kD Mini-PROTEAN TGX Precast Protein Gels (Bio-Rad), SDS-PAGE gels, and PVDF membranes were used for the separation and transfer of proteins, respectively. Primary antibodies and HRP conjugated secondary antibodies used in this study are listed in supplemental Table 3. Chemiluminescence detection was captured using the ChemiDoc Imaging System with Image Lab Software (Bio-Rad).

**IL-1 treatment**

For immunoblotting studies, leukemia KG-1a, THP-1, and U937 cells were serum-starved overnight in base medium plus 0.5% FBS. Subsequently, human IL-1β was added to the cells (KG-1a: 2 ng/mL; THP-1 and U937: 10 ng/mL) and incubated for 0, 5, 10, 30, 60, and 120 minutes. Primary human AML cells and NBM cells were cultured in cytokine SFM, followed by stimulation with human IL-1β (10 ng/mL) for 0 and 5 minutes. At each time point, cells were collected, washed with cold 1x PBS, and lysed for immunoblotting assays.

#### Supplemental Figures

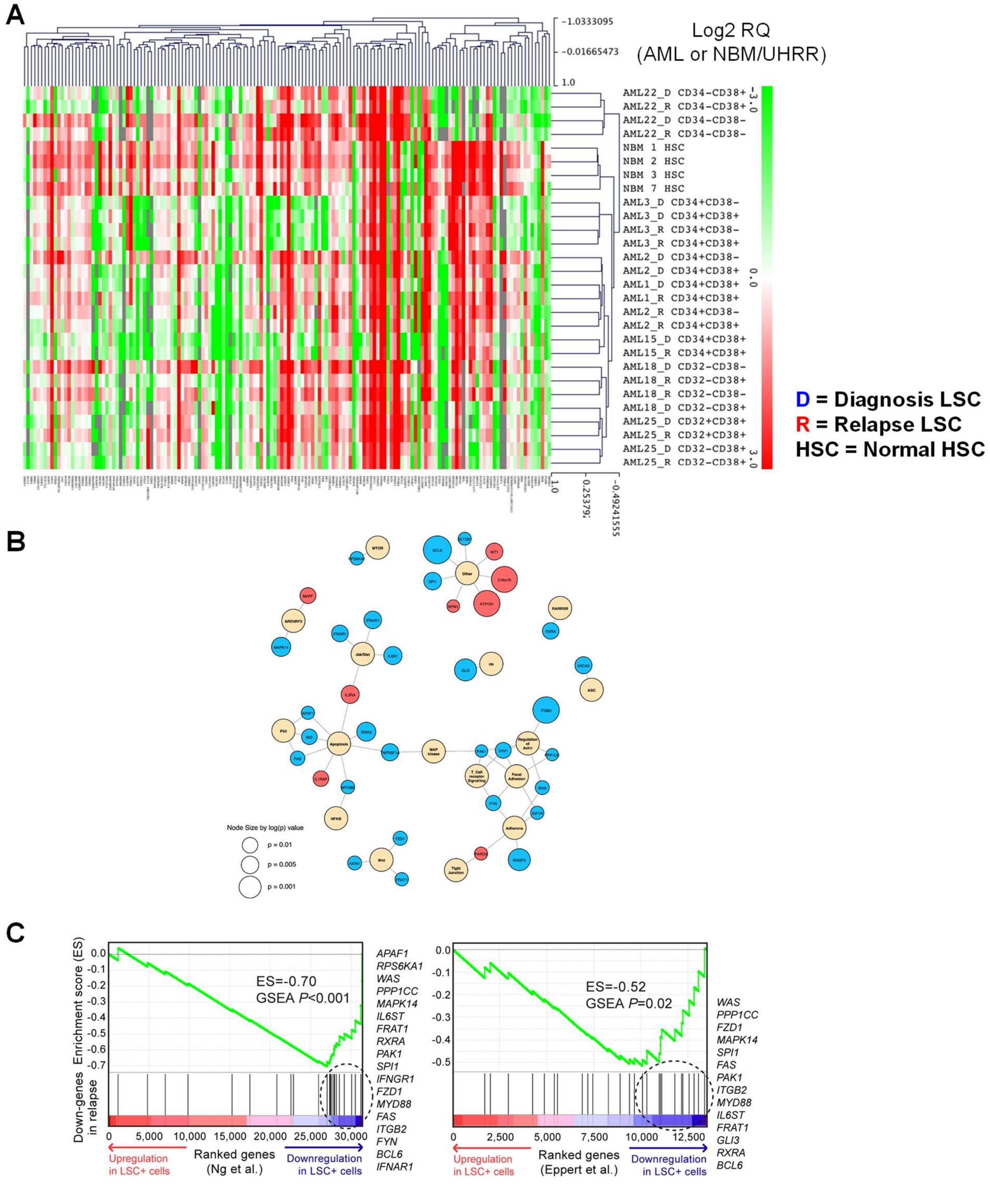


**Figure S1. Targeted transcriptome analyses of paired AML diagnostic and relapsed LSC populations and normal HSC populations.**

(A) Heat map displaying the normalized relative expression of LSC-related genes in paired AML diagnostic and relapsed LSC populations, as well as normal HSC populations. Twelve paired AML LSC populations ($CD3^-CD34/CD38$ or $CD3^-CD32/CD38$) from seven samples and four NBM HSC populations ($CD34^+CD38^-CD90^+Lin^-$) were analyzed using TLDA-based analyses. Data were clustered using Pearson correlation metric with average linkage. RQ = Relative Quantification. UHRR = Universal Human Reference RNA. WT1 is not included in this heat map. (B) Network plots depicting the signaling pathway interactions among a total of 36 dysregulated genes (DEGs) identified in AML relapsed LSCs (diagnosis vs. relapse; paired t-test $P < 0.05$). Among these DEGs, 8 are upregulated (indicated in red), and 28 are downregulated (indicated in blue). Each DEG is connected to the related pathway (indicated in yellow), and the node size represents the log($P$) value. (C) GSEA plots comparing genes downregulated upon relapse with published expressional profiles of $LSC^+$ cells. Genes were ranked based on the significance of differential expression associated with $LSC^+$ cells (indicated by red and indigo arrows) of two published datasets, *Eppert et al.* (6) and *Ng et al*. (7). Twenty-eight downregulated genes were defined as a gene set (black segments) and tested for the enrichment (enrichment score; green curve) on either side of the ranked list. Denoted by a dashed circle are leading-edge component genes that contribute to the enrichment. Significance was assessed using the random permutation test.

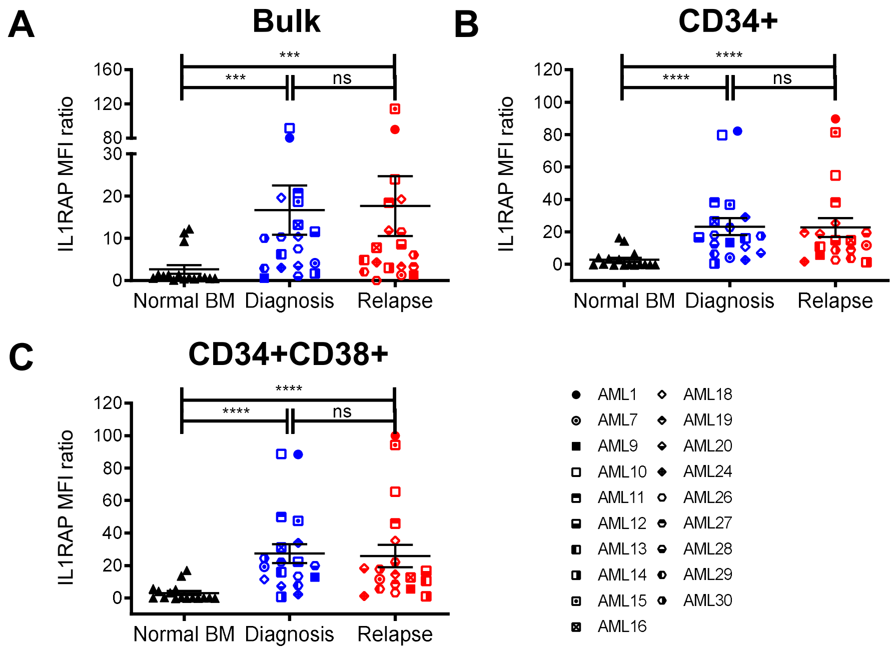


**Figure S2. Immunophenotyping of IL1RAP on primary AML samples at diagnosis and relapse compared to NBM counterparts.**

Protein expression levels of IL1RAP in bulk (A), CD34$^{+}$ (B), and CD34$^{+}$CD38$^{+}$ (C) populations on primary AML samples at diagnosis and relapse, as well as NBM counterparts (n = 16). Median fluorescence intensity (MFI) of the cell-surface IL1RAP was quantified by flow cytometry analysis and normalized to Fluorescence Minus One (FMO) control. MFI ratio = (Sample MFI – FMO MFI)/FMO MFI. Mean ± standard error of the mean (SEM). The two-tailed Mann-Whitney test (NBM vs. AML) or Wilcoxon matched-pairs signed rank test (diagnosis vs. relapse) were used in comparison. ***$P \leq 0.001$, ****$P \leq 0.0001$, ns = not significant.

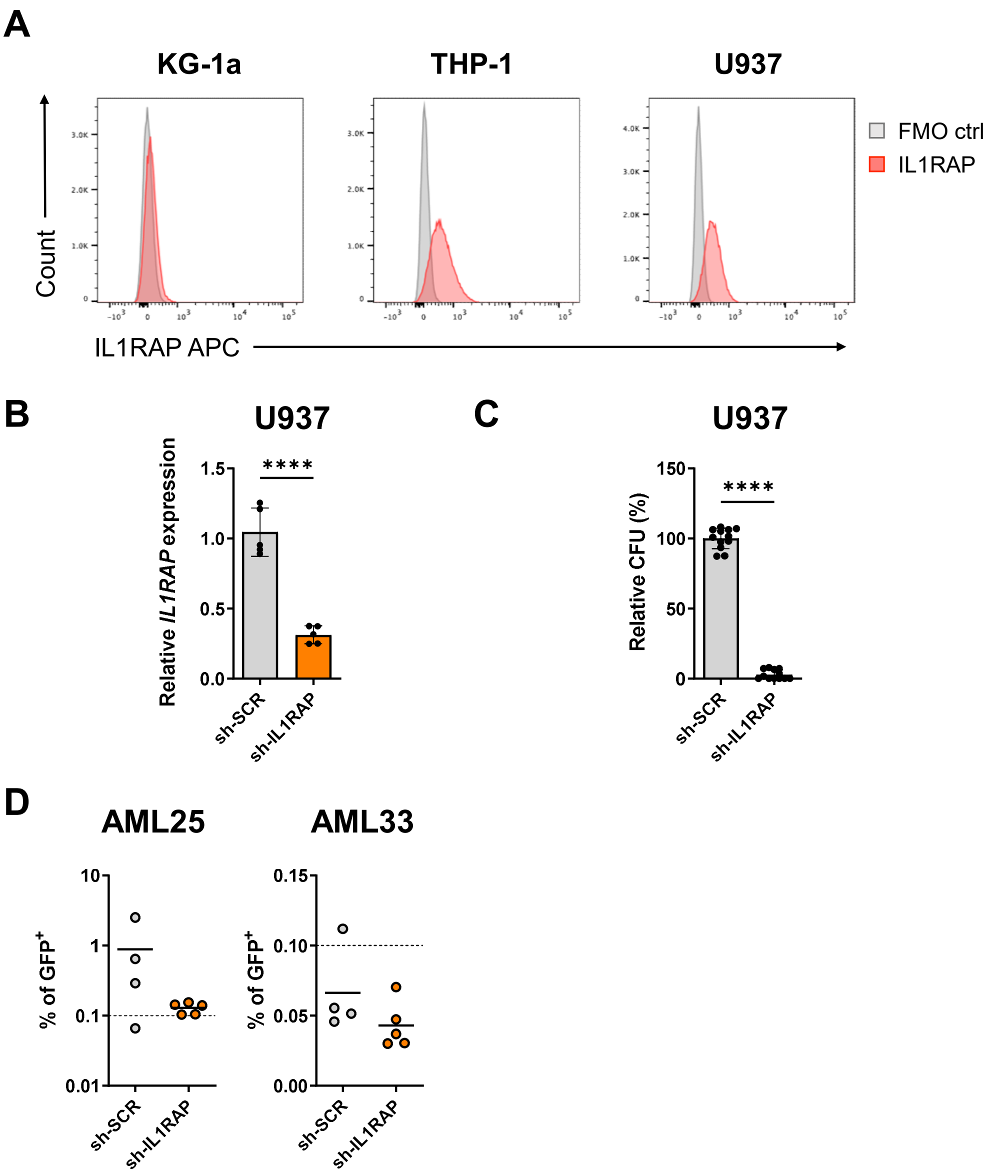


**Figure S3. Loss of *IL1RAP* affects LSC function in human leukemia cells.**
(A) Representative protein expression profiles of IL1RAP in human leukemia cell lines KG-1a, THP-1, and U937 analyzed using flow cytometry. n = 3 independent experiments. (B) RT-qPCR analyses of U937 cells expressing IL1RAP shRNA (sh-IL1RAP) and scramble shRNA (sh-SCR). Total RNA was isolated from sorted live $GFP^+$ cells at 4 days post infection. n = 5 independent experiments. Mean ± SD. Unpaired t test. (C) CFU-C assays of U937 cells expressing sh-IL1RAP and sh-SCR. Sorted live $GFP^+$ cells were plated 3 days post infection, and the CFU-C was counted after 14 days of culture. n = 4 independent experiments. Mean ± SD. Unpaired t test. (D) Xenotransplantation assays of primary AML cells (AML25 and AML33) with or without knockdown of IL1RAP. Each dot represents the $GFP^+$ engraftment level in the BM of individual NSG mouse analyzed by flow cytometry analysis. The horizontal line represents the mean of $GFP^+$ engraftment level in NSG mice of each group. Unpaired t test. ****$P \leq 0.0001$.

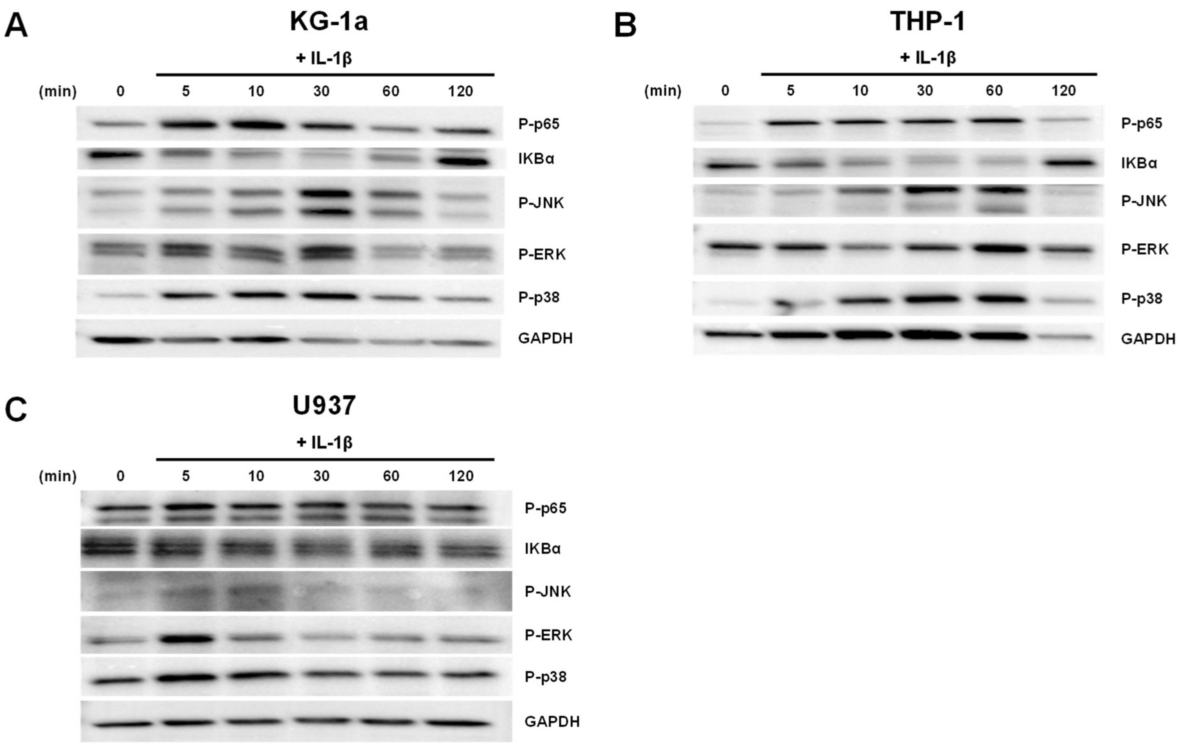


**Figure S4. Immunoblotting analyses of IL-1 signaling key factors in human leukemia cells.** Leukemia cell lines KG-1a (A), THP-1 (B) and U937 (C) were serum-starved overnight in base medium plus 0.5% FBS. Subsequently, human IL-1β was added to the cells (KG-1a: 2 ng/mL; THP-1 and U937: 10 ng/mL) and incubated for 0, 5, 10, 30, 60, and 120 minutes. Cells were collected and lysed at each time point for immunoblotting analyses.

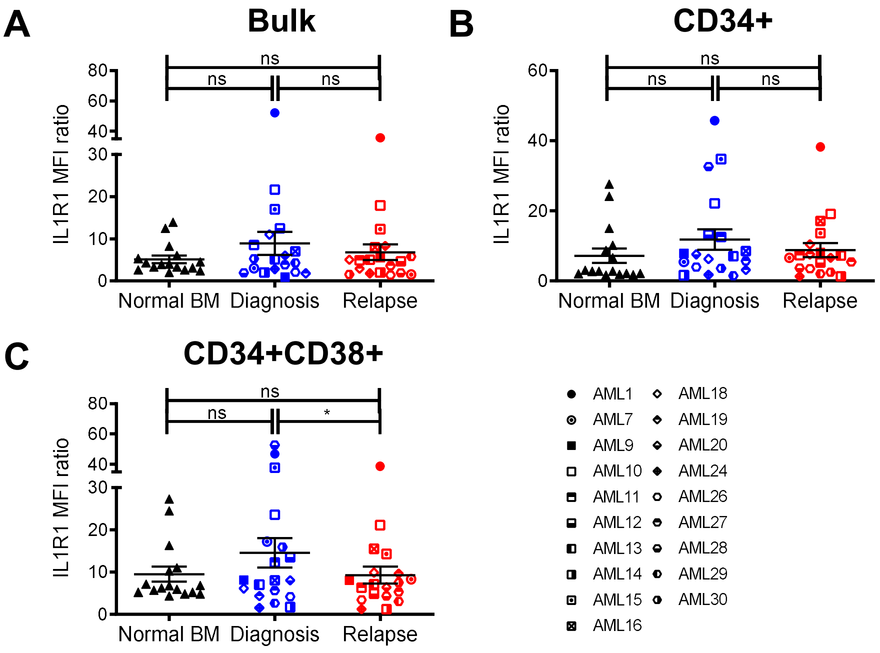


**Figure S5. Immunophenotyping of IL1R1 on primary AML samples at diagnosis and relapse compared to NBM counterparts.**

Protein expression levels of IL1R1 in bulk (A), $CD34^+$ (B), and $CD34^+CD38^+$ (C) populations on primary AML samples at diagnosis and relapse, as well as NBM counterparts (n = 16). Median fluorescence intensity (MFI) of the cell-surface IL1RAP was quantified by flow cytometry analysis and normalized to Fluorescence Minus One (FMO) control. MFI ratio = (Sample MFI – FMO MFI)/FMO MFI. Mean ± SEM. The two-tailed Mann-Whitney test (NBM vs. AML) or Wilcoxon matched-pairs signed rank test (diagnosis vs. relapse) were used in comparison. *$P \leq 0.05$, ns = not significant.

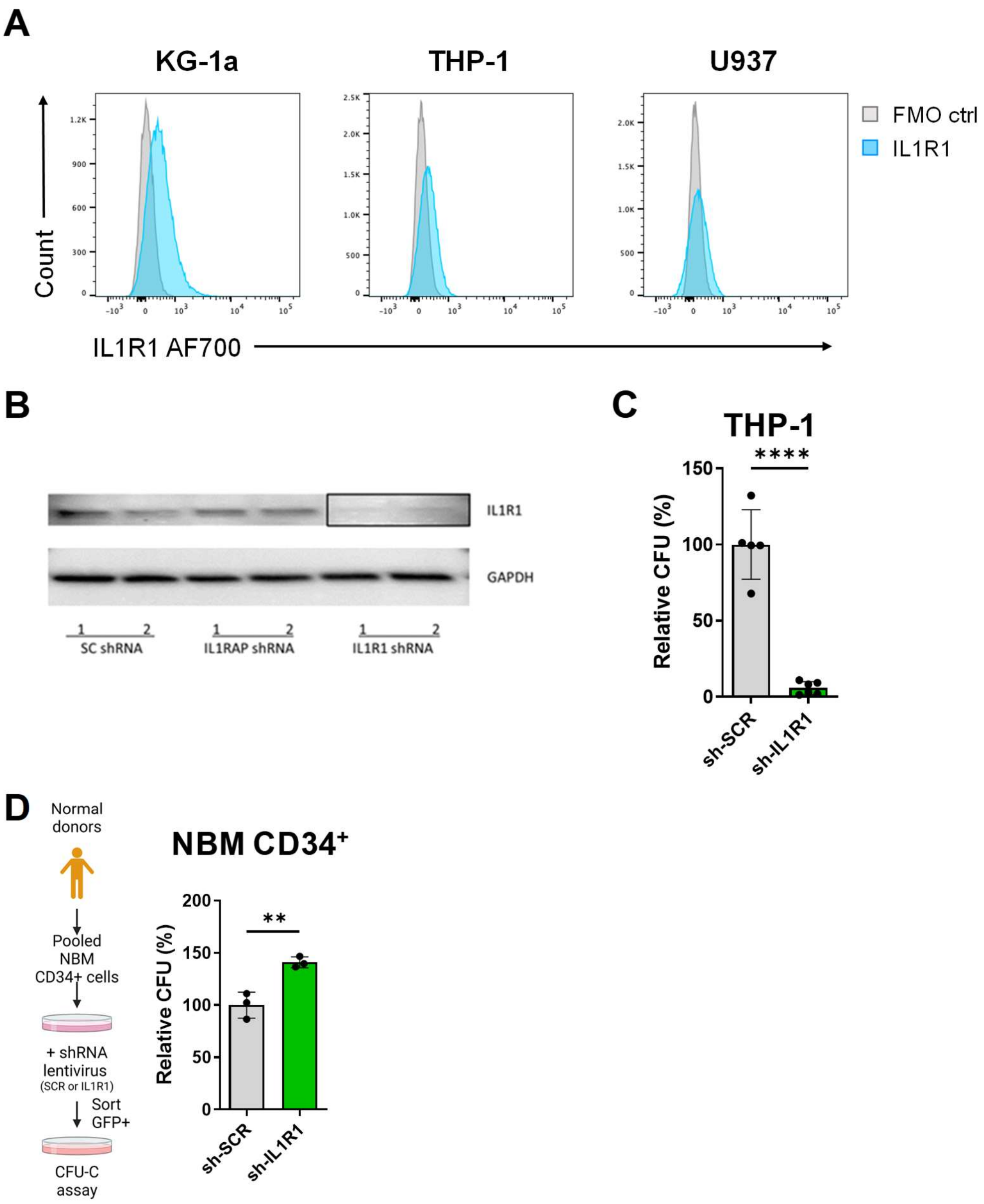


**Figure S6. Loss of *IL1R1* affects LSC function in AML cells.**
(A) Representative protein expression profiles of IL1R1 in human leukemia cell lines KG-1a, THP-1, and U937 analyzed using flow cytometry. n = 3 independent experiments. (B) Protein expression levels of IL1R1 in human AML BM endothelial cells expressing sh-IL1R1, sh-IL1RAP, and sh-SCR. After 3-4 days of infection, cells were treated with or without IL-1β (2 ng/mL) for 5 minutes and harvested for immunoblotting assays. 1 = Untreated. 2 = Treated with IL-1β. (C) CFU-C assays of THP-1 cells expressing sh-IL1R1 and sh-SCR. Cells were plated for CFU-C at 3 days post infection. n = 2 independent experiments. Mean ± SD. Unpaired t test. (D) CFU-C assays of NBM CD34$^+$ cells with or without knockdown of IL1R1. Mean ± SD. Unpaired t test. $^{**}P \leq 0.01$, $^{****}P \leq 0.0001$.

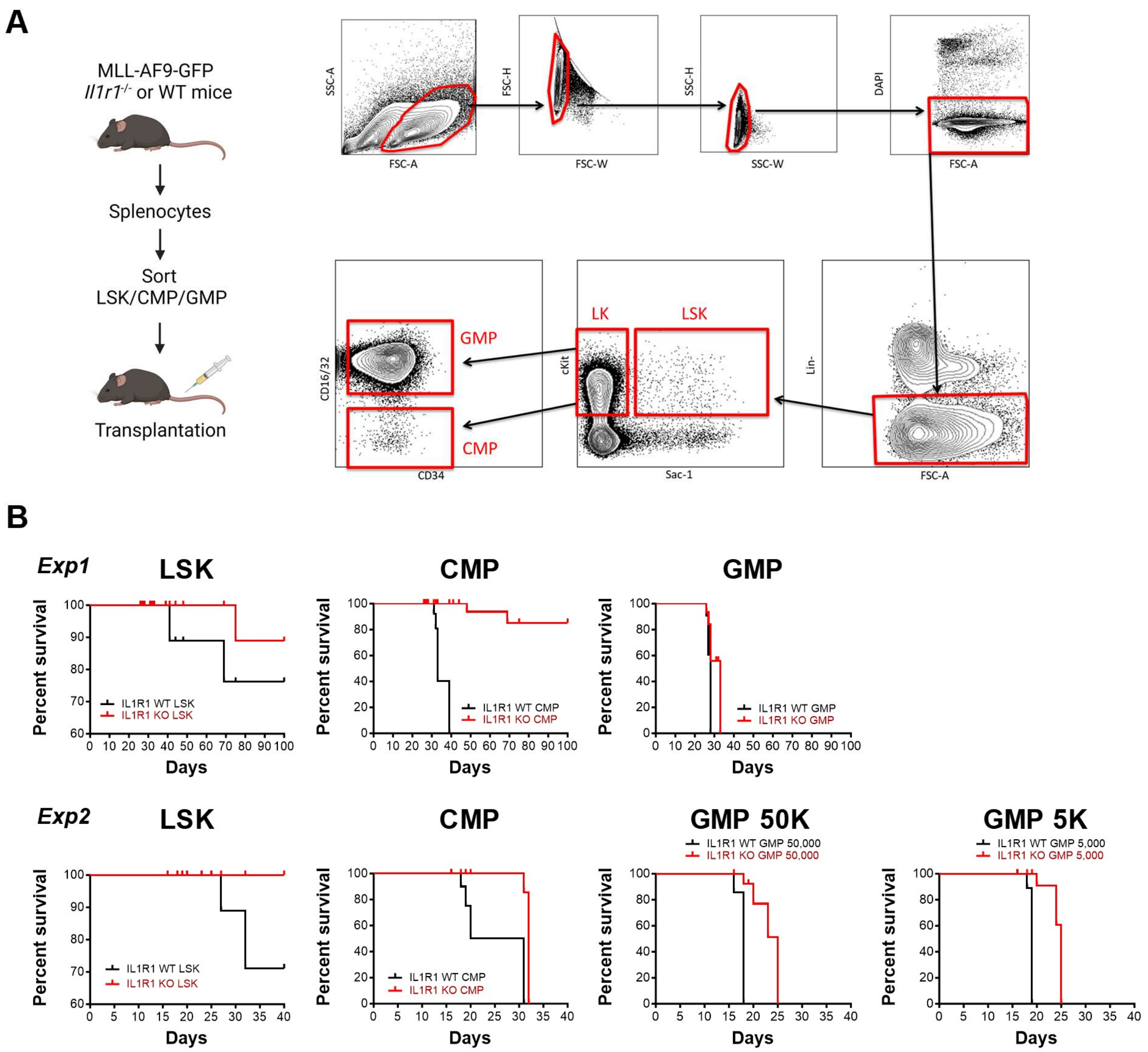


**Figure S7. Loss of IL1R1 impairs the engraftment activity of leukemia LSK, CMP, and GMP populations.**

(A) Schematic of the experiment strategy for transplantation using MLL-AF9 LSK, CMP, and GMP populations with or without IL1R1 knockout. In brief, IL1R1 WT and IL1R1 KO MLL-AF9-GFP splenocytes from primary recipients were sorted for leukemia LSK, CMP, and GMP populations (LSK: $Lin^{-}Sca\text{-}1^{+}c\text{-}Kit^{+}$; CMP: $Lin^{-}c\text{-}Kit^{+}(16/32)FC\gamma R^{-}CD34^{+}$; GMP: $Lin^{-}c\text{-}Kit^{+}(16/32)FC\gamma R^{+}CD34^{+}$) and then transplanted into secondary Pep Boy recipients. Mice were closely monitored until the time of death. (B) Overall survival analyses of recipients transplanted with MLL-AF9 LSK, CMP, and GMP populations, either with or without IL1R1 knockout, as described in *(A)*. The figures display results from two independent studies (*Exp 1* and *Exp 2*). The cell count for each population was determined based on the proportion of cells identified during sorting. The smallest population served as the reference. GMP 50K and GMP 5K represent 50,000 and 5,000 GMP cells per mouse, respectively. Schematics were created with BioRender.com.

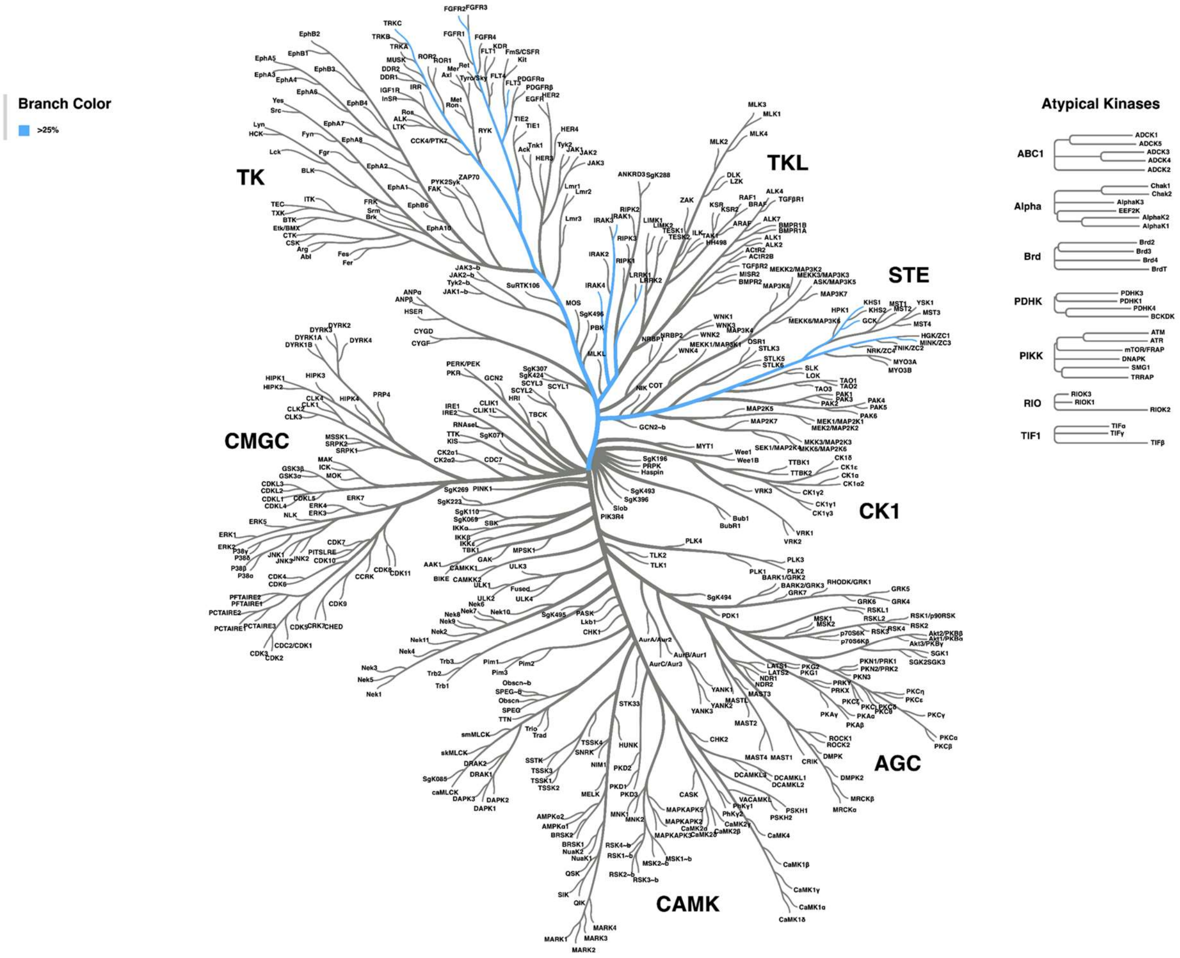


**Figure S8. Kinome map analysis of UR241-2.**

The kinome tree map of UR241-2 was created using Coral generated from the Phanstiel Lab. Blue lines highlight kinases that exhibit inhibition above the cut-off threshold of 25%, affected by UR241-2 at a single dose of 5 nM. TK = Tyrosine Kinase. TKL = Tyrosine Kinase-Like kinases. STE = Homologs of yeast Sterile 7, Sterile 11, and Sterile 20 kinases. CK1 = Casein Kinase 1. AGC = PKA, PKG, and PKC families. CAMK = Calcium/calmodulin-dependent protein kinase. CMGC = CDK, MAPK, GSK3, and CLK families.

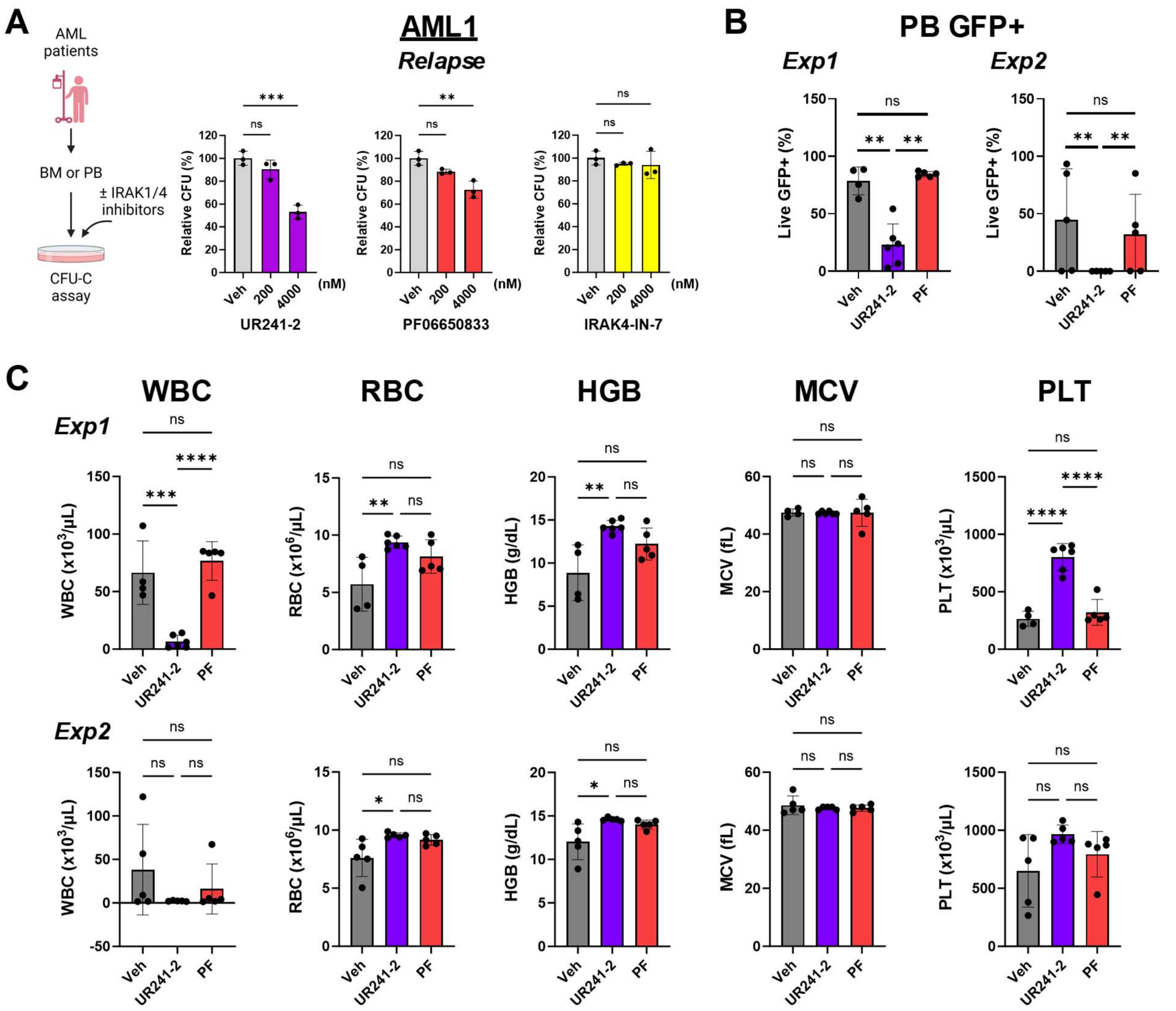


**Figure S9. UR241-2 suppresses LSC function and engraftment activity in AML.**
(A) CFU-C assays of relapsed primary human AML cells (AML1). Cells were treated with IRAK1/4 inhibitors UR241-2, PF06650833, and IRAK4-IN-7 (0.2 and 4 μM) in methylcellulose-based medium. Mean ± SD. Ordinary one-way ANOVA followed by Dunnett's multiple comparisons test. (B) Engraftment levels of *ex vivo* treated MLL-AF9 GFP$^+$ cells in the peripheral blood (PB) in recipients at four weeks after transplantation, as described in *Figure 5D*. Mean ± SD. Two-tailed Mann-Whitney test. (C) Complete blood count (CBC) values in the PB in recipients, received *ex vivo* treated MLL-AF9 GFP$^+$ cells, at four weeks after transplantation as described in *Figure 5D*. Mean ± SD. Ordinary one-way ANOVA followed by Tukey's multiple comparisons test. *$P \leq 0.05$, **$P \leq 0.01$, ***$P \leq 0.001$, ****$P \leq 0.0001$, ns = not significant. Schematics were created with BioRender.com.

#### Supplemental Tables

| Identifier | Sex | Age (y) | Cytogenetics | BM Blasts (%) | WBC (K) | Mutations | Induction | RFS (m) |
|---|---|---|---|---|---|---|---|---|
| **AML1** | M | 58 | Normal | 60-65 | 80 | ND | FLAG | 5.2 |
| **AML2** | F | 70 | Normal | 44 | 18.4 | ND | Ida + Bortezomib | 10.3 |
| **AML3** | M | 67 | t(3;16), del(5)(q?31), -9, -17, -18, +2mar | 74 | 13.1 | FLT3, p53 | 7+3 | 4.4 |
| **AML7** | F | 39 | Normal | 60 | 10.9 | N/A | 7+3 | 14.4 |
| **AML9** | F | 23 | +1, der(1;15)(q10;q10) | 83 | 5.2 | N/A | 7+3 | 8.3 |
| **AML10** | F | 57 | Normal | 81 | 171.6 | DNMT3a, FLT3, NPM1 | 7+3 | 6.2 |
| **AML11** | N/A | 69 | +8, inv(9)(p21q21)?c | 91 | 54.5 | DNMT3a, FLT3 | ADE | 12.1 |
| **AML12** | N/A | 23 | inv(16)(p13q22) | 56 | 92.2 | NRAS | HiDAC/Ida | 3.7 |
| **AML13** | N/A | 71 | del(20)(q11q13) | 90 | 118.1 | RUNX1, EZH2 | HiDAC/Ida | 20.5 |
| **AML14** | N/A | 72 | Normal | 48 | 11.4 | IDH1 | ADE | 6.8 |
| **AML15** | N/A | 68 | Normal | 39 | 3.7 | DNMT3a, FLT3, NPM1, NRAS | ADE | 9.0 |
| **AML16** | N/A | 78 | t(9;22)(q34;q11) | 54 | 77.4 | ND | ADE | 7.4 |
| **AML18** | N/A | 80 | Normal | 94 | 68.5 | DNMT3a, NPM1, TET2 | ADE | 13.3 |
| **AML19** | N/A | 65 | +11 | 77 | 77.7 | RUNX1, FLT3 | HiDAC/Ida | 8.6 |
| **AML20** | N/A | 76 | Normal | 28 | 3.2 | IDH2, ASXL1, | 7+3 | 17.3 |
| **AML22** | N/A | 60 | +8 | 80 | 29.5 | IDH1, NRAS, TET2 | 7+3 + Zosuquidar | 15.5 |
| **AML24** | N/A | 73 | +2mar | 29 | 49.7 | DNMT3a, TET2 | ADE | 14.0 |
| **AML25** | M | 50 | 45, X, -Y, t(9;11)(p22;q23) | 87 | 95.8 | FLT3 | 7+3 | 7.2 |
| **AML26** | M | 57 | trisomy 11 | 78 | 4.4 | ND | 7+3 | 48 |
| **AML27** | F | 41 | 46 (X;X) | 61 | 13.6 | FLT3 | 7+3 | 9 |
| **AML28** | F | 55 | trisomy 21 | 15 | 2.8 | ND | DEC/7+3 | 6 |
| **AML29** | F | 66 | Normal | 28 | 61.6 | ND | CLO | 3 |
| **AML30** | M | 61 | Normal | 60 | 125.5 | FLT3ITD, NPM1 | 7+3 | 10 |
| **AML31** | N/A | N/A | N/A | N/A | N/A | N/A | N/A | N/A |
| **AML33** | F | 71 | Normal | 87 | 178.9 | FLT3ITD, NPM1 | 7+3 | N/A |
| **AML34** | M | 50 | Normal | 99 | 113.4 | FLT3ITD | 7+3/5+2 | N/A |
| **AML35** | N/A | N/A | Normal | N/A | 100 | ND | N/A | N/A |

**Table S1. Characteristics of AML specimens.**
The clinical information for AML1 to AML25 is adapted from *Ho et. al.*'s study (1). Additionally, we have included nine other AML specimens for immunophenotyping or

functional assays, as listed in this table. RFS = relapse-free survival. ND = not detected. N/A = data not available.

| Gene List of TLDA | | | | | | | |
|---|---|---|---|---|---|---|---|
| ABCA5 | CCNB1 | EIF2S3* | IL3RA | MYD88 | PPP2R1B | RIPK1 | TNFRSF1A |
| ABCB1 | CCND2 | EVI1 | IL6ST | NAB1** | PPP2R5C | RPS6 | TRAF3IP2 |
| ABCG1 | CCND3 | FAS | INSR | NCKAP1 | PPP3CB | RPS6KA1 | UBR5* |
| ACTG1 | CCNG2 | FER | IQGAP1 | NCOA1 | PPP3CC | RXRA | ULK2 |
| ACTN4 | CDH1 | FLJ13197* | IQGAP2 | NCOA2 | PRICKLE1 | SERPINE1 | VAV1 |
| AKT3 | CDH2 | FLNB | IRAK3 | NCRNA00338;SEC14L1* | PRKAR1A | SESN3 | VEGFA |
| APAF1 | CDK12** | FRAT1 | ITGA4 | NF1** | PRKAR2B | SETDB1** | VGLL4** |
| APC2 | CHEK1 | FRMD4B* | ITGB2 | NFATC1 | PRKCB1 | SMAD2 | WAS |
| ARFGEF1** | CLN5* | FYN | ITGB7 | NFATC2 | PTEN | SMAD3 | WASF1 |
| ARHGEF12 | CREB1 | FZD1 | JUN | NIPAL2* | PTK2 | SPI1 | WASF3 |
| ATP1B1* | CSDE1* | FZD2 | LRRC8B* | NPM1 | PTPN6 | SRF | WIF1 |
| AXIN1 | CSNK1A1 | FZD6 | MAFF | PAK1 | PTPRM | SSH3 | WT1 |
| BAD | CSNK1G1 | GLI3 | MAFG | PARD3 | RABGAP1** | SSX2IP | ZFP30* |
| BCL2 | CSNK2A1 | GPR56* | MAP3K7* | PLCB4 | RAC1 | STAT6 | ZMAT3 |
| BCL2L1 | CTBP2 | HMGB3 | MAPK14 | PMAIP1 | RAPGEF2 | STRADA | 18S |
| BCL6 | CYCS | IFNAR1 | MAPK8 | PNPLA4** | RARA | SUFU | GAPDH |
| BID | DIAPH1 | IFNGR1 | MEIS1 | POLR2A | RASGRP1 | TCF7L2 | HPRT1 |
| BIRC3 | DUSP10 | IGF1R | MKNK1 | PPIG* | RBPMS** | TGIF2** | |
| C16orf5* | DVL3 | IKBKG | MPL | PPP1CC | RFWD2 | TNFAIP3* | |
| CACYBP | EGR1 | IL1RAP | MYB | PPP1R10* | RHEB | TNFRSF10B | |

**Table S2. The 157 gene list of TaqMan Low Density Array (TLDA) cards for targeted transcriptome analysis.**

Our previous study identified 3,005 differentially expressed genes (DEGs) and 10 dysregulated pathways in enriched phenotypically defined LSC populations ($CD34^{+}CD38^{-}CD123^{+}Lin^{-}$ or $CD34^{+}CD38^{-}CD90^{-}Lin^{-}$) compared to normal HSC populations ($CD34^{+}CD38^{-}CD90^{+}Lin^{-}$) using genome-wide expression analyses and two independent pathways analyses (8). In this study, we selected 124 genes from the 3005 DEGs based on their involvement in five dysregulated pathways (Adherens Junction, Wnt, Regulation of Actin Cytoskeleton, Jak-STAT, and MAPK) or reported roles in response to therapy. Additionally, we included 27 genes (labeled as *) from a previously reported 42 gene LSC-related (LSC-R) gene signature, which compared expression profiles of functionally defined LSC populations to non-LSC populations, regardless of surface antigen phenotype (6). Notably, 10 of these 27 genes overlapped with our 3,005 DEG list (labeled as **). Furthermore, we incorporated 3 additional cancer-related genes (BCL6, CDH1, and EGR1), along with 3 reference genes (18S, GAPDH, and HPRT1).

**<u>Flow cytometry analysis:</u>**

**Anti-human antibodies**

| Target | Conjugate | Clone | Brand |
|---|---|---|---|
| CD3 | Biotin | UCHT1 | BD Pharmingen |
| CD3 | PE-Cy5, PE-Cy7 | HIT3a | BioLegend |
| CD14 | Biotin | 61D3 | eBioScience |
| CD15 | Biotin | HI98 | eBioScience |
| CD16 | Biotin | 3G8 | BD Pharmingen |
| CD19 | Biotin, PE | HIB19 | BD Pharmingen |
| CD32 | APC, PE-Cy7 | 6C4 | eBioScience |
| CD34 | BUV395 | 581 | BD Horizon |
| CD34 | FITC | 581 | BD Pharmingen |
| CD34 | PE-Cy7 | 8G12 | BD |
| CD38 | APC | HB7 | BD |
| CD38 | PerCp-Cy5.5 | HIT2 | BD Pharmingen |
| CD45 | PE-Cy5 | HI30 | BioLegend |
| CD56 | Biotin | B159 | BD Pharmingen |
| CD90 | APC | 5E10 | BD Pharmingen |
| CD97 | BV605 | VIM3b | BD OptiBuild |
| CD97 | FITC | VIM3b | BD Pharmingen |
| CD123 | PE | 7G3 | BD Pharmingen |
| CD123 | PE-CF594 | 7G3 | BD Horizon |
| Glycophorin A | PE-Cy5 | GA-R2 | BD Pharmingen |
| IL1R1 | AF700, PE | 129304 | R&D Systems |
| IL1RAP | APC, PE | 89412 | R&D Systems |
| Streptavidin | PE-Cy5 | -- | BD Pharmingen |

**Anti-mouse antibodies**

| Target | Conjugate | Clone | Brand |
|---|---|---|---|
| B220 | Biotin | RA3-6B2 | eBioscience |
| CD3e | Biotin | 145-2C11 | BD Pharmingen |
| CD16/32 | APC-Cy7 | 2.4G2 | BD Pharmingen |
| CD34 | PE | RAM34 | BD Pharmingen |
| CD45 | APC | 30-F11 | BD Pharmingen |
| CD45.1 | APC-Cy7 | A20 | BD Pharmingen |
| CD45.2 | APC-Cy7 | 104 | BioLegend |
| c-Kit | PE-Cy5 | 2B8 | eBioscience |
| GR-1 | Biotin | RB6-8C5 | eBioscience |
| Sca-1 | PE-Cy7 | D7 | BD Pharmingen |
| Ter119 | Biotin | Ter-119 | eBioscience |
| Streptavidin | PE-CF594 | -- | BD Horizon |

**Table S3. Antibodies used in this study.**

**Immunoblotting:**

| Antibodies | Cat# | Brand |
|---|---|---|
| Phospho-NF-κB p65 (Ser536) | #3033 | Cell Signaling |
| Phospho-p38 MAPK (Thr180/Tyr182) | #9211 | Cell Signaling |
| Phospho-Akt (Ser473) (D9E) XP | #4060 | Cell Signaling |
| Phospho-p44/42 MAPK (Erk1/2) (Thr202/Tyr204) (D13.14.4E) XP | #4370 | Cell Signaling |
| NF-κB p65 (D14E12) XP | #8242 | Cell Signaling |
| p38 MAPK | #9212 | Cell Signaling |
| Akt | #9272 | Cell Signaling |
| p44/42 MAPK (Erk1/2) (137F5) | #4695 | Cell Signaling |
| IL1R1 | #MAB269 | R&D Systems |
| IL-1β (3A6) | #12242 | Cell Signaling |
| α-Tubulin (DM1A) | #3873 | Cell Signaling |
| GAPDH (14C10) | #2118 | Cell Signaling |
| Anti-rabbit IgG, HRP-linked | #7074 | Cell Signaling |
| Anti-mouse IgG, HRP-linked | #7076 | Cell Signaling |

**Table S3 (cont'd). Antibodies used in this study.**

#### Supplemental References


1. Ho TC, LaMere M, Stevens BM, Ashton JM, Myers JR, O'Dwyer KM, et al. Evolution of acute myelogenous leukemia stem cell properties after treatment and progression. *Blood.* 2016;128(13):1671-8.
2. Ashton JM, Balys M, Neering SJ, Hassane DC, Cowley G, Root DE, et al. Gene sets identified with oncogene cooperativity analysis regulate in vivo growth and survival of leukemia stem cells. *Cell Stem Cell.* 2012;11(3):359-72.
3. Neering SJ, Bushnell T, Sozer S, Ashton J, Rossi RM, Wang PY, et al. Leukemia stem cells in a genetically defined murine model of blast-crisis CML. *Blood.* 2007;110(7):2578-85.
4. Ye H, Adane B, Khan N, Alexeev E, Nusbacher N, Minhajuddin M, et al. Subversion of Systemic Glucose Metabolism as a Mechanism to Support the Growth of Leukemia Cells. *Cancer Cell.* 2018;34(4):659-73 e6.
5. Ye H, Adane B, Khan N, Sullivan T, Minhajuddin M, Gasparetto M, et al. Leukemic Stem Cells Evade Chemotherapy by Metabolic Adaptation to an Adipose Tissue Niche. *Cell Stem Cell.* 2016;19(1):23-37.
6. Eppert K, Takenaka K, Lechman ER, Waldron L, Nilsson B, van Galen P, et al. Stem cell gene expression programs influence clinical outcome in human leukemia. *Nat Med.* 2011;17(9):1086-93.
7. Ng SW, Mitchell A, Kennedy JA, Chen WC, McLeod J, Ibrahimova N, et al. A 17-gene stemness score for rapid determination of risk in acute leukaemia. *Nature.* 2016;540(7633):433-7.
8. Majeti R, Becker MW, Tian Q, Lee TL, Yan X, Liu R, et al. Dysregulated gene expression networks in human acute myelogenous leukemia stem cells. *Proc Natl Acad Sci U S A.* 2009;106(9):3396-401.
